# Pattern formation along signaling gradients driven by active droplet behaviour of cell groups

**DOI:** 10.1101/2024.04.08.588511

**Authors:** Hugh Z Ford, Giulia L Celora, Elizabeth R Westbrook, Mohit P Dalwadi, Benjamin J Walker, Hella Baumann, Cornelis J. Weijer, Philip Pearce, Jonathan R Chubb

**Author notes:** Co-first authors (Hugh Z Ford, Giulia L Celora).

## Abstract

Gradients of extracellular signals organise cells in tissues. Although there are several models for how gradients can pattern cell behaviour, it is not clear how cells react to gradients when the population is undergoing 3D morphogenesis, in which cell-cell and cell-signal interactions are continually changing. *Dictyostelium* cells follow gradients of their nutritional source to feed and maintain their undifferentiated state. Using light sheet imaging to simultaneously monitor signaling, single cell and population dynamics, we show that the cells migrate towards nutritional gradients in swarms. As swarms advance, they deposit clumps of cells at the rear, triggering differentiation. Clump deposition is explained by a physical model in which cell swarms behave as active droplets: cells proliferate within the swarm, with clump shedding occurring at a critical population size, at which cells at the rear no longer perceive the gradient and are not retained by the emergent surface tension of the swarm. The droplet model predicts vortex motion of the cells within the swarm emerging from the local transfer of propulsion forces, a prediction validated by 3D tracking of single cells. This active fluid behaviour reveals a developmental mechanism we term “musical chairs” decision-making, in which the decision to proliferate or differentiate is determined by the position of a cell within the group as it bifurcates.

## Introduction

Signaling gradients are interpreted by cells to guide their migration and to direct the sub-division of embryonic tissues into specific cell types (*1, 2*). Despite the widespread functioning of gradients in both development and disease, it has remained challenging to monitor natural signaling gradients together with cell and tissue responses over time. In contexts with limited tissue reorganisation, it has been possible to infer how cells react to signal gradients (*3*). However, for contexts in which three-dimensional tissue organization remodels substantially over time, there are significant barriers to interpreting the connection between signal inputs and behavioural outputs of cells. In these systems, the organization of cells continually changes, influencing and being influenced by cell-cell interactions (*4, 5*) and extracellular signal gradients (*6-11*) in addition to any emergent tissue properties, which all combine to influence the cell response to signaling.

In this study, we investigate the emergent dynamics and organization of cell groups migrating towards self-generated signaling gradients. We use light sheet imaging to simultaneously monitor the dynamics of a nutritional signaling gradient and its effects on the migration and differentiation of populations of *Dictyostelium* cells. We show how the gradient organises single cells into dense groups - swarms. These swarms periodically shed large cell clumps, driving the cells in the clumps into the developmental programme. Clump shedding is surprising in the light of traditional models of collective cell chemotaxis along self-generated gradients (*12, 13*), so to explain this emergent behaviour, we developed and tested a coarse-grained mathematical model in which the cell swarm is represented as an active droplet. Our model implies that an emergent surface tension is a key determinant of pattern formation in these chemotactic cell populations. The model also predicts an emergent vortex motion of cells within the swarm, which our experiments confirm is a key driver of cell transport. Behaviours of the cell swarm arising from droplet properties (shedding and vortex motion) combine to determine cell fate: the position of the cell in the vortex at the time of clump shedding dictates whether or not the cell enters the developmental programme.

## Results

### Splitting of cell swarms during chemotaxis

*Dictyostelium* cells use signaling gradients to coordinate their differentiation programme. In their undifferentiated proliferative state, these soil-dwelling amoebae locate their nutritional source, bacteria, by chemotaxis towards bacterial metabolites (*14*). Without bacteria, the cells starve and enter their developmental programme, in which single cells form multicellular aggregates via chemotaxis towards cAMP, before forming a final structure carrying dormant spores. To mimic natural environments, we spotted cells on lawns of their bacterial food source (*15-17*). Macrophotography shows the proliferating cell population clearing the bacterial lawn as an advancing ring-shaped band, called the feeding front (Figure 1A, Video 1). Compact clumps are shed from the feeding front, which collectively form a spotted pattern (Supplementary Figure 1A). These clumps emerge from patches along the advancing feeding front that elongate as finger-like protrusions, then pinch off and settle into isolated domes (Figure 1A’, Supplementary Figure 1C, Video 2). Further from the feeding front are the first clear signs of development: cell streaming and aggregation, characteristic of cAMP chemotaxis (Figure 1A 2^nd^ panel and 1A’ last panel).

**Figure 1.**
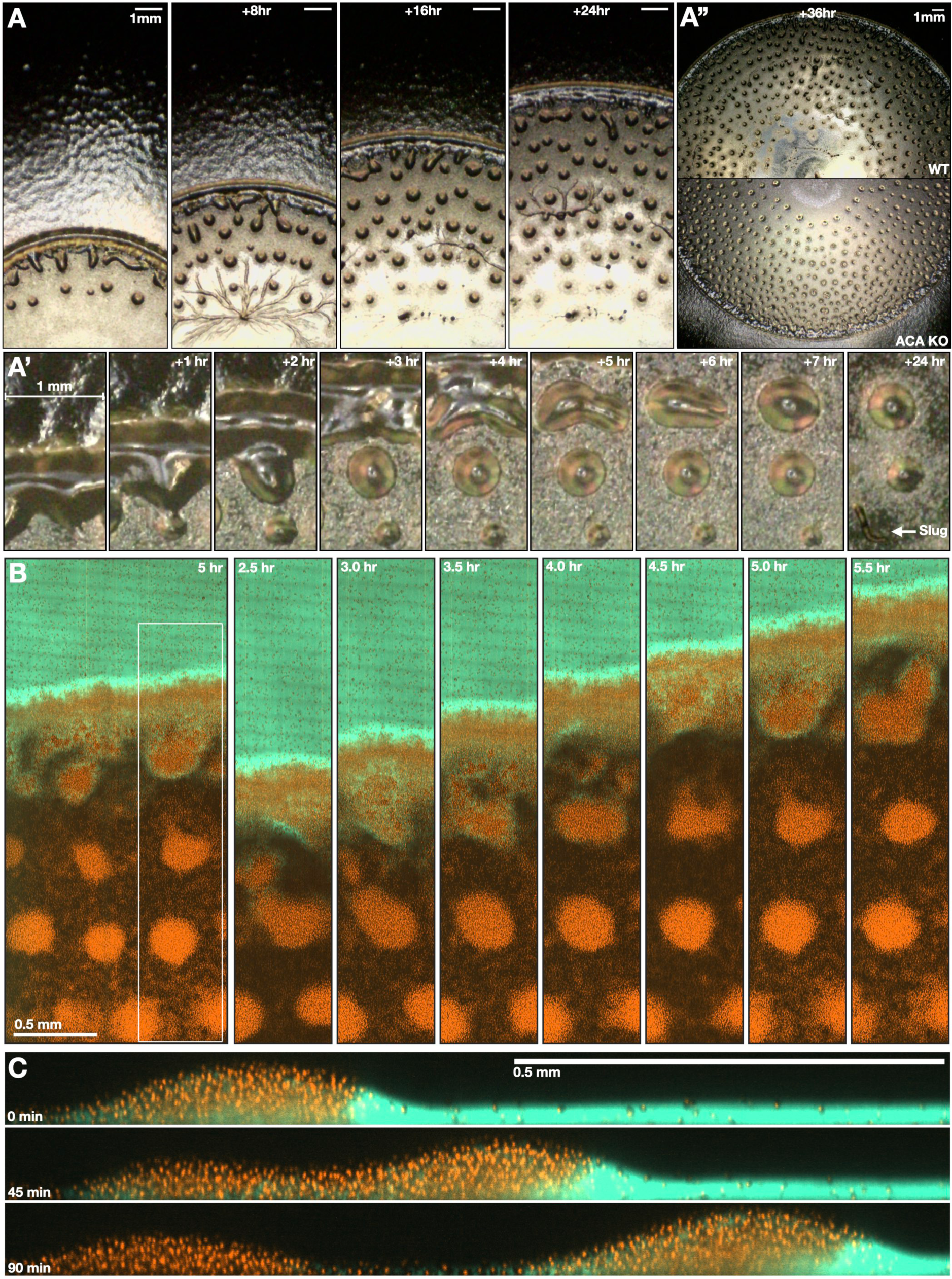
Multi-scale imaging of cell migration to signaling gradients. **A** Macrophotography of the feeding front, see also Video 1. *Dictyostelium* cells progressively clear the bacteria as an expanding ring-shaped band. Cells are left behind (inside the ring) as isolated cells and compact cell clumps. **A’** Close-ups of 2 cell shedding events. **A’’** A comparison to feeding fronts of *acaA*-cells (no cAMP signaling; top right). Scale bars: 1mm. **B** Light sheet imaging of feeding fronts. 3D images of fluorescently labelled bacteria (green) and *Dictyostelium* cell nuclei (red) at the feeding front, see also Video 3. Scale bar: 500µm. The right and central panels show the progression of the feeding front and clump formation within the region marked by the white box in the left panel. **C** Similar to B, but showing a side-view of the feeding front at 15 minute intervals, see also Video 4. Scale bar: 500µm.

To investigate the cell-cell and cell-signal interactions at the feeding front requires distinguishing between *Dictyostelium* cells and the bacteria. For this we used light sheet microscopy to live image fluorescently-labelled bacteria and cell nuclei at the millimetre scale over multiple hours (Figure 1B & 1C, Supplementary Figure 2A-2C, Video 3 & 4). These data show that the cells advance at the interface with the bacteria as a densely packed and highly motile swarm. This behaviour is characteristic of classic Keller-Segel models of chemotaxis (*12, 13*) where motile cell groups self-generate a chemoattractant gradient and remain at a constant size due to a balance of cell growth and continuous cell shedding (*18, 19*). However, as implied by the macrophotography (Figure 1A), and in contrast to predictions from classic Keller-Segel models, in addition to continuous cell shedding, most cells (∼60-70%) are left behind in large, stable, and spatially compact clumps that are distinct from the field of isolated cells (Figure 1B & 1C, Supplementary Figure 3E). The cell clumps do not reengage with the advancing front, indicating that the cells within them are destined to starve and then enter the developmental programme.

Multicellular development in *Dictyostelium* is dependent on the chemoattractant cAMP. To test whether the shedding of cell clumps requires cAMP, we analysed clump shedding in cells lacking *acaA*, the gene encoding the adenylyl cyclase synthesising cAMP during starvation (Figure 1A’’, Supplementary Figure 1B, Video 1). In *acaA-* cells, clump shedding occurs with the same characteristics as wild-type, indicating shedding does not require cAMP. Indeed, without cAMP, clumps are abnormally persistent, implying cAMP is required for clump dispersal, not formation. Cells within clumps withstand cAMP signaling for multiple cycles of streaming and aggregation before they disperse; clump dispersal is necessary for the transition to the multicellular structures of late stages of the developmental programme (Figure 1A’, Supplementary Figure 1C, Video 1 & 2). Consequently, development is suspended by 1-2 days for the cells in clumps compared to cells outside clumps. This spontaneous heterogeneity in developmental timing may provide flexibility within the population to counter uncertain nutrient availability or variance in the opportunity to disperse spores.

### Clump shedding follows gradient dynamics

Based on this initial analysis, we infer that shedding of clumps from the feeding front emerges from physical interactions between cells and/or interactions between cells and the bacterial gradient. To determine how swarm motion and clump shedding relate to the gradient, we quantified the dynamics of swarm size together with the distribution of bacteria (Figure 2A, Supplementary Figure 2B & 3A, Video 3 & 5). Based on Keller-Segel models, cells feeding on a bacterial lawn should generate a stable exponential or logistic decay in the quantity of bacteria across the swarm (*12, 13*). However, our data show the bacterial gradient is highly dynamic and can flatten and even reverse towards the rear of the swarm, corresponding to a minimum gradient of less than zero within the swarm boundary (Figure 2B & 2C, Supplementary Figure 4A, 4B & 5B). Additionally, as observed previously (*15*), bacteria accumulate along the feeding front, creating a local peak with up to twice the quantity of bacteria ahead of the swarm (Figure 2B, Supplementary Figure 5A). To understand the basis of the bacteria peak, we used particle image velocimetry (PIV) to quantify the bulk motion and interactions of the swarm and bacteria (Supplementary Figure 6A & 6B, Video 4). This analysis implies the bacteria are pushed forward, with the swarm acting analogous to a snowplough, suggesting the bacterial population possesses a material integrity that provides resistance to swarm penetration (Supplementary Figure 6C). The bacteria peak remains a constant size at the swarm front consistent with a balance between accumulation via swarm motion and degradation via feeding. This persistent bacteria accumulation creates a robust, positive bacteria gradient localised at the swarm front, potentially allowing long-range cell migration for cells at the leading edge, but not necessarily the rear, as the position of the negative gradient varies with respect to the rear of the swarm (Figure 2B & 2D, Supplementary Figure 4B & 5B). Overall, these results imply the swarm shapes the chemoattractant gradient by both spatially reorganising and degrading the bacteria.

**Figure 2.**
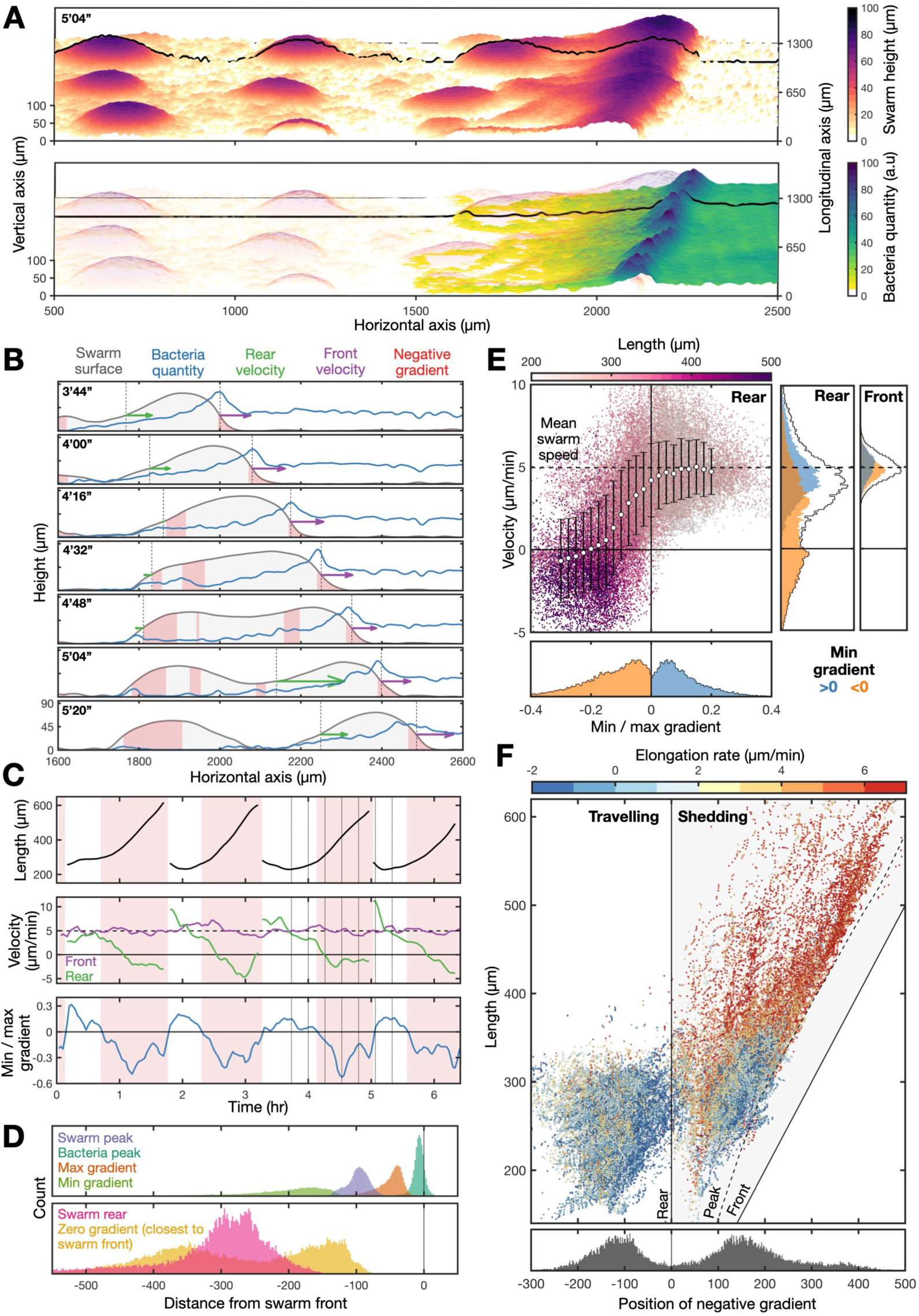
Coupled swarm and signaling gradient dynamics trigger collective cell shedding. **A** Quantification of the cell swarm height and bacteria quantity at the feeding front (from data in Fig 1B). Black lines mark the swarm boundary (top panel) and bacterial quantity (bottom panel) at one cross section. **B** Swarm shape dynamics during clump shedding. Time lapse (h’min’’) of the cross sections of the swarm height (grey) and bacteria quantity (blue) shown in A throughout one shedding event. Also shown are: the regions where the gradient is negative (red), swarm front and rear (black dotted lines), and velocity vectors of the swarm front (purple) and rear (green). **C** Tracking the dynamics of swarm length (black) and bacterial gradient (blue) for a cross-section of the feeding front, through several shedding cycles. Shedding events are defined as a collapse in swarm length. Swarm elongation is summarised by diffferences in velocity of the swarm front (purple) and rear (green). The bacterial gradient is summarised as the ratio of the minimum gradient and maximum gradient within the swarm. Times where the minimum gradient is negative are highlighted by red blocks. Vertical lines in the third shedding cycle correspond to sequence of plots in B. The mean swarm speed (4.9 µm/min) is shown as a black dashed line. **D** The bacterial peak always coincides with the swarm front. Plots summarise gradient and swarm properties at all time and spatial points. Top plot shows distances between the locations of swarm front (black line) and the: swarm and bacterial peak, and the maximum and minimum gradient within the swarm. Bottom plot shows how the zero gradient can be positioned either side of the swarm rear. **E** Elongation (reduction of rear velocity) only occurs when the minimum gradient becomes negative within the swarm. Main plot shows the relationship between the rear velocity, length (colour) and gradient (ratio of the minimum and maximum gradient). Bottom panel shows the gradient is partitioned into positive and negative values. Side panels show the front speed is independent of the minimum gradient, whereas the rear speed changes from the mean swarm speed to around zero when the minimum gradient becomes negative. **F** Two phases of swarm behaviour: travelling and shedding. Top panel shows two clusters of swarm behaviour differing in the position of the negative gradient (minimum distance from the swarm front) relative to the swarm rear, swarm length and elongation rate (colour), for all spatial and time points. Bottom plot: two discrete clusters showing locations of the zero gradient is either behind (left) or deep within the swarm boundary (right).

Tracking the dynamics of swarm size and bacterial gradient for a cross-section of the feeding front, through several shedding cycles (Figure 2C, Supplementary Figure 4B & 4C), reveals how clump shedding is preceded by: i) steady swarm elongation, ii) a reduction of the rear velocity of the swarm to around zero and iii) the emergence of a negative gradient within the swarm boundary. Indeed, comparing these three quantities across whole datasets shows that the swarm rear becomes stationary only once the minimum gradient within the swarm is less than zero (Figure 2E). Incorporating spatial information (tracking the position of the furthest forward negative gradient) into this analysis reveals two clusters corresponding to two phases of swarm dynamics: travelling and shedding (Figure 2F). The position of the negative gradient relative to the swarm rear is defined such that it is positive when it is within the swarm, and negative when behind. The travelling phase is characterised by a compact swarm (length below 350µm) with a positive minimum gradient (Figure 2F, Supplementary Figure 5D). In this phase, the swarm size remains stable because cells at the rear move at a similar speed to cells at the front, causing no swarm elongation (Figure 2E, Supplementary Figure 5C). This is consistent with all cells in the swarm having access to a positive bacterial gradient and therefore adequate positional information on the location of the bacterial food source. In contrast, the shedding phase is characterised by a steady swarm elongation rate (around the mean swarm speed) and a negative gradient deep within the swarm boundary (Figure 2E & 2F, Supplementary Figure 5D). This elongation is consistent with cells at the swarm rear lacking positional information derived from the bacterial front.

What causes the loss of positional information within the swarm and why does this cause collective shedding? As might be expected due to cell growth and proliferation, our data show a steady and low baseline rate of swarm elongation during the travelling phase (Supplementary Figure 5C). Swarms above a critical length then rapidly elongate and eventually split (Figure 2F, Supplementary Figure 5C & 5E), suggesting cells at the swarm rear lose positional information because they are too far from the bacterial source. However, this interpretation does not account for the bulk shedding of cell clumps, because continuous cell growth would steadily push cells beyond the critical length at the rear, resulting in a continuous shedding of cells. Therefore, loss of positional information caused by growth does not seem to be sufficient to explain collective cell shedding. Alternatively, redistribution of bacteria across the swarm (Supplementary Figure 6C, Video 4) could cause a sudden emergence of a negative gradient associated loss of positional information from the front. However, swarms initially maintain compactness beyond the first emergence of a negative gradient at the swarm rear (Figure 2F). Indeed, even during splitting events, the swarm maintains a smooth and consistent boundary until it pinches off (Figure 2B, Supplementary Figure 4D & 5D). This suggests that some emergent material property of the multicellular swarm combines with growth and loss of positional information to trigger the transition to shedding.

### Active fluid model of cell swarming

Based on the high density of cells and the clearly delineated swarm boundary observed in the experiments (Figure 1A, Supplementary Figure 1A), we hypothesised that the swarm has emergent fluid-like properties caused by physical interactions between cells. That is, we interpret the cell swarm as a living active droplet (*20-23*). To investigate whether and how emergent fluid-like properties determine the observed swarm dynamics, we developed a continuum, coarse-grained, active fluid thin-film model for a cross-section of the swarm (Figure 3A, Supplementary Information – Mathematical Modelling).

**Figure 3.**
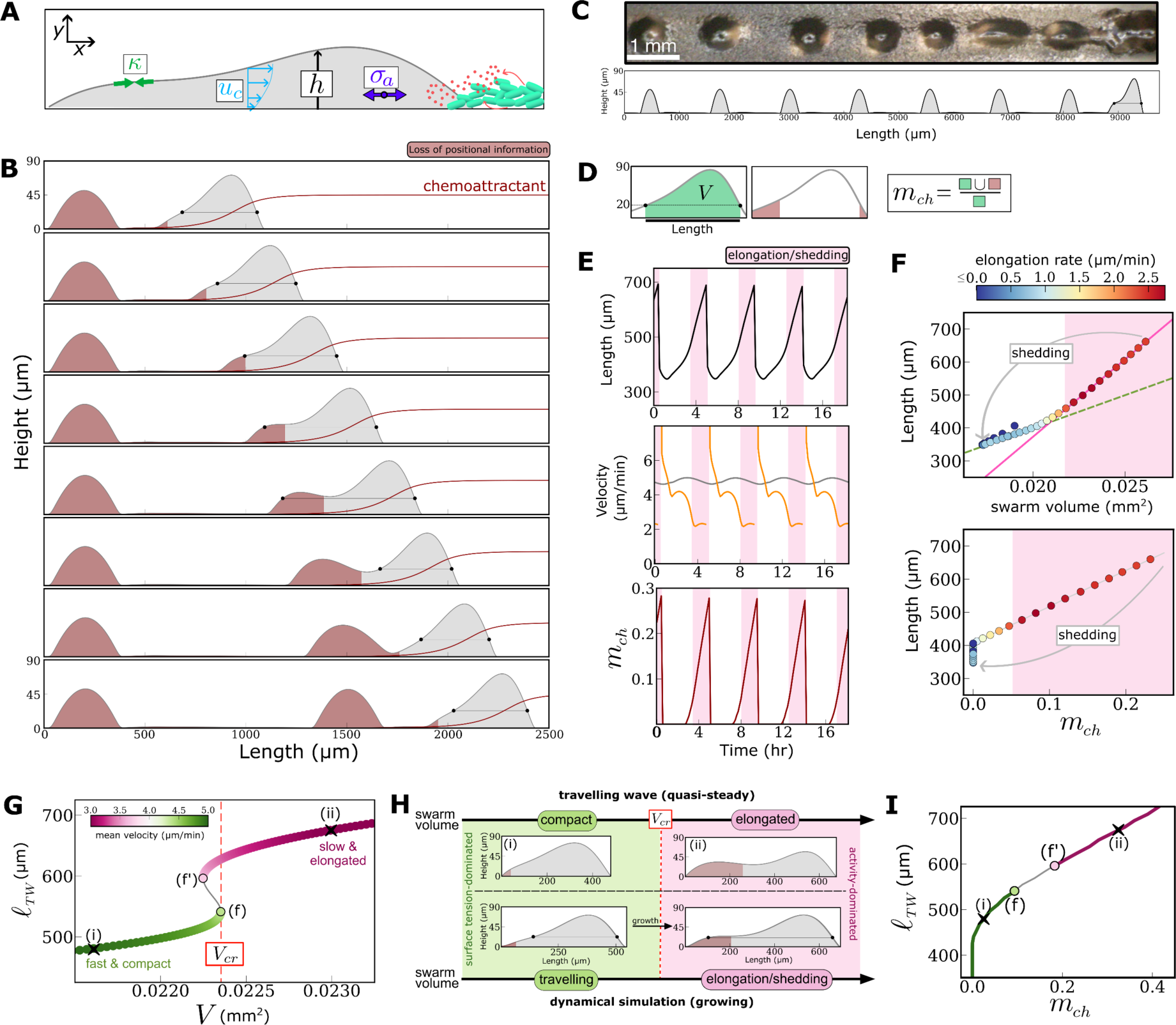
Active Fluid Model of Cell Swarms. **A** Schematic summarising the key features of the active fluid model (Modelling Supplement). **B** Time-lapse (25 minutes intervals) of the simulation of the active fluid thin film model. We plot the swarm height (grey curve) and the normalised concentration of chemoattractant (dark red curve). Also shown in dark red is the region where the gradient in the chemoattractant drops below a cell sensitivity threshold (Modelling Supplement). The black dots show the points at which the swarm height drops below a 20 *μm* threshold (Figure 3D). **C** Comparison between 1D-track experiment (top figure; viewed from above) and model simulations of a swarm cross section (bottom figure). The model captures the periodic shedding of clumps and quantitatively matches their distance (1.25 mm) and shedding rate (4.6 hours). **D** Schematic summarising how we extract swarm length, swarm volume, *V*, and the metric, *m_ch_*, from the output of the dynamic simulation of the active fluid model. **E** The length (black), velocity of the swarm front (grey) and rear (orange), and the ratio of the swarm mass exposed to a chemoattractant gradient below the cell sensitivity threshold (*m_ch_*, dark red), throughout 4 shedding cycles. **F** Two-phase (travelling and elongation/shedding) of the swarm dynamics shown via the relationship between swarm length and normalised swarm mass (top plot) and the ratio of the swarm mass exposed to a chemoattractant gradient below the cell sensitivity threshold *m_ch_* (bottom plot). Transition to swarm elongation is signified by a rapidly increasing length relative to mass (top panel) and the key swarm quantity *m_ch_*increasing above a critical (non-zero) threshold. In both plots the colour indicates the elongation rate. **G** Bifurcation diagram of travelling wave solution for the active fluid droplet model illustrating how the length of the swarm (coloured by the swarm TW velocity) changes as a function of the swarm mass. Bifurcation points (f & f’) delimit a region of bistability. The presence of the two fold points introduces a discontinuous transition in the mode of swarm migration which is controlled by the volume increasing above the critical value, *V_cr_*. **H** Schematic illustrating the connection between the travelling wave analysis and the two phase regime observed in the dynamical simulations. Top plots showing the height profile for the travelling wave solutions corresponding to point (i) and (ii) in the bifurcation diagram (Figure 3F). Bottom plots showing the height profile for the swarm front in the dynamical simulations (extracted from first and third panels in Figure 3B). **I** Two-types of TW swarm solution shown via the relationship between the length of the swarm and the metric *m_ch_*.

In the model, we assumed that the following three material properties contribute to emergent fluid stress in the swarm: i) an effective surface tension *κ*, which generates capillary stresses that confine the advancing cell swarm, and give a propensity for circularity and consistent contact angles in stationary cell clumps (Supplementary Figure 3B-3D); ii) an effective viscosity *η* which generates viscous stresses; this property arises from the turnover of cell-cell attachments and rearrangements of cells within the swarm (*24*); iii) an activity parameter *ξ*, associated with an effective active contribution σ_*a*_ to the stress in the fluid, which arises from the alignment of directed cell motion due to chemotactic bias. Cell proliferation was modelled through film growth, which is mediated by the local concentration of bacteria. In the model, bacteria are consumed by cells, and produce diffusible chemoattractant molecules that decay at a constant rate. To capture the emergent flow of cells inside the swarm, we used lubrication theory, which is valid for long, thin films representing the swarms studied here (Figure 1 & 2). We applied a Navier slip condition at the floor to account for effective friction with the floor (*25*). Under these assumptions, the flow of cells in the swarm, *u*_*c*_, has a parabolic profile (see Figure 3A); the magnitude of the flow depends on the relative sizes of surface tension, viscosity and active stress gradients. The model suggests that the emergent flow field causes the swarm to migrate up self-generated chemoattractant gradients, which are in turn shaped by feeding (Figure 3B).

To calibrate the model, we estimated the emergent material properties of the swarm (Supplementary Information – Mathematical Modelling), by quantitatively matching model predictions for the shedding rate and distance between shed clumps to experimental data (Figure 3C & Supplementary Information – Mathematical Modelling). For this purpose, we used macrophotography to live image feeding fronts constrained to thin lines of bacteria, which enables unambiguous measurement of the shedding rate (Supplementary Figure 7A-7C). In this context, we find that the shedding is periodic with rate of 1 clump per 4.35 hours, which matches the proliferation rate of cells feeding on bacteria (*26*). Calibrated model simulations recapitulate the two observed phases of the swarm movement: travelling and shedding (Figure 3B, 3E & 3F, Video 6). During the travelling phase, the swarm rear and front move at a similar velocity (Figure 3D & 3E) and during the shedding phase, the model predicts a rapid increase in swarm length due to the rear of the droplet moving slower than the front, in agreement with experimental observations (Figure 1C, 2E). Elongation terminates with a shedding event, after which the swarm re-enters the travelling phase. Because proliferation continues in the swarm front after shedding, this results in periodic shedding events, which are clearly observed when cell swarms migrate along 1D lines of bacteria (Figure 3C and Supplementary Figure 7A). We conclude that our minimal model can explain the emergent swarm dynamics observed experimentally.

We then explored the physical reasons for the observed periodic shedding dynamics. The elongation phase in both model and data corresponds to an increase in the ratio between the elongation rate and swarm expansion (Figure S6B, Figure 3F top panel). This suggests that the rapid elongation of the swarm before shedding is not caused by a sudden increase in proliferation, but rather by a redistribution of the mass within the swarm. To determine how the swarm transitions from compact propagation to elongation and shedding, we considered a simplified version of our model, in which the swarm is described as a travelling droplet with quasi-constant volume *V*. In this simplified framework, we find that compact, fast-travelling swarms can only exist for swarm volumes *V* below a critical value *V*_)*_ (SI Text, Figure 3G & 3H). For swarm volumes above this critical value, the model predicts only elongated, slow-travelling swarms (Figure 3G). For increasing values of *V*, the compact swarm solution transitions to the elongated solution across a discontinuous phase transition (mathematically a fold bifurcation (*27*)) at *V* = *V*_)*_. Physically, the transition is explained by the competition between capillary forces generated by surface tension, which favour swarm compactness, and chemotaxis-driven gradients in active stress, which favour the elongation of the swarm. For larger swarms, active stress gradients are increased owing to the finite chemoattractant decay length. This explains why the elongated solution is exhibited above a critical swarm volume. In the dynamical simulations, the critical volume is reached because of slow cell proliferation; then, the crossing of the bifurcation forces the swarm to reassemble over a timescale much faster than growth, eventually leading to shedding (Figure 3H). The model predicts that the critical volume depends on the material properties – an increase in surface tension widens the surface tension-dominated regime, and can suppress shedding entirely (Modelling Supplement Figures 5 and 7).

Because chemotaxis-driven gradients in active stress drive shedding in the dynamical simulations, we quantified their effect by introducing the swarm metric, *m_ch_*, which measures the proportion of the swarm mass with insufficient positional information for directed motion, i.e. an (almost) flat chemoattractant gradient (Figure 3D). In the simulations, an increase of this metric above a critical value coincides with the transition between the surface tension-dominated (compact) and activity-dominated (elongated) regimes (Figure 3F & 3I). Experimentally, loss of positional information can be approximated with the flattening/reversal of the bacterial gradient. When restructuring experimental data (Figure 2E) by *m_ch_*, we find experimental curves that are consistent with model predictions (Supplementary Figure 5E). Overall, our model and experiments suggest that the swarm behaves as an active droplet that migrates in self-generated chemotactic gradients: the transition between travelling and shedding phases is a proliferation-driven transition between a surface tension-dominated regime (compact swarm) and an activity-dominated regime (larger swarms elongating and shedding). More broadly, our results indicate that emergent material properties of physically interacting cells can drive pattern formation in response to signaling gradients.

### Implications of swarm active fluid behaviour for individual cells

A key prediction of the active droplet model of cell swarming is the form of the emergent vortex flow-field in the frame of reference of the swarm (Figure 4A & 4B, Supplementary Information). To experimentally test this form of collective cell motion, we tracked the 3D positions of individual cells from high spatial and temporal resolution live imaging of swarms (Supplementary Figure 8A-8C, Video 7). To study the internal flow of cells, we averaged cell velocities across the face of the swarm (Figure 4C, Supplementary Figure 9A-9C). This analysis shows that cell motion within the swarm is indeed spatially organised as a vortex, where cells at the top of the swarm move faster than cells at the bottom (Figure 4D). During the travelling phase of swarm migration, conservation of mass is achieved by cells at the swarm rear climbing to the surface and cells at the swarm front moving to the floor, with a cycle time of around 2 hours along the outer swarm boundary (Figure 4B & 4D). In both the model and data, the location where the cells are moving at the mean group speed aligns tightly with the half-swarm height (Figure 4A & 4C). During the shedding phase, both model and data indicate a stable vortex cell flow that persists throughout swarm elongation, resulting in a larger contribution to clumps from cells at the swarm floor (Figure 4E).

**Figure 4.**
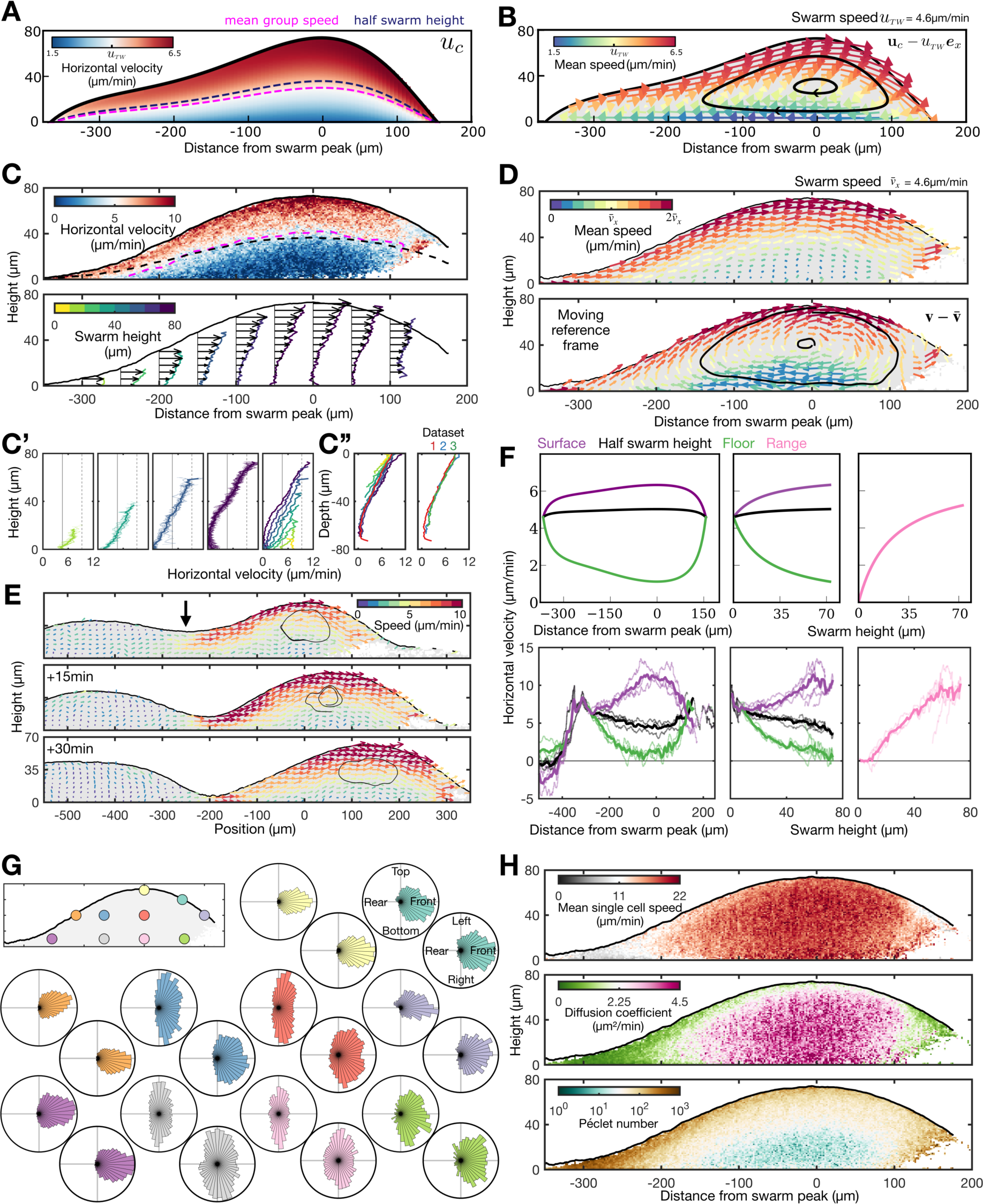
Validating the cell flow patterns predicted by the active fluid model. **A** Model predictions for the horizontal velocity field within the swarm, labelled with contours for mean group speed and half swarm height. **B** Model predictions for the vector field of cell flows in the moving reference frame of the swarm, labelled with streamlines summarising cell circulation. **C** Mean cell horizontal velocity field obtained from cell tracks. Plots also show contours for the half-swarm height (black) and mean group speed (pink). Bottom panel shows vector field of horizontal velocities, summarised with curves coloured by swarm height. **C’** shows these curves partitioned into different heights, with the far right panel combining these mean horizontal velocity profiles for different swarm heights. In contrast, in **C’’** these curves are structured by swarm depth rather than height. Left panel is from one dataset, right panel shows averaged curves from 3 datasets. **D** Validation of model predictions showing vortex flow in the swarm. Plots derived from 3D cell tracking data and plotted as in B. **E** Vortex fields derived from 3D cell tracking data during a shedding event. **F** Comparing cell velocity profiles between model (top panels) and data (bottom panels). Left panels: mean horizontal cell velocity at the surface, floor and half swarm height at different distances to the swarm peak. Middle panels: horizontal velocities as a function of swarm height. Right panels show the range (distance between surface and floor velocities). **G** Polar distributions of cell orientations at various points across the face of the swarm (inset panel). Each position (colour) has two plots showing the 2D planes of the side and top views. **H** The distribution of the mean speed of individual cells (top), the diffusion coefficient (middle) and the Peclet number (bottom).

What are the causes and consequences of the emergent vortex flow-field? Our data show that differences in cell motion along the vertical axes have a stronger relationship with distance from the swarm surface (depth) than from the floor (height), indicating that vortex motion does not solely arise from a boost in motion by cells riding on those below them (Figure 4C’ & 4C’’). Instead, the model predicts that cells at the top of the swarm move faster than cells at the bottom because of an effective friction with the floor which, combined with viscosity, leads to a Poiseuille-like flow profile (Figure 4A-4C) (*28*). In the model, viscosity arises from the dynamic attachments between cells, and causes cell motion to be dependent on the motion of surrounding cells – cells exert forces on their neighbours to propel themselves. Consequently, the overall flow of cells is more directed at the top of the swarm than at the floor because directed cell motion near the floor is reduced by the effective friction, which in turn reduces the directed motion of neighbouring cells. In other words, the model predicts that the emergent material properties of the swarm suppress the ability for cells at the swarm floor to respond to the gradient. This effect is clearly apparent in our data showing the differences in cell motion at each horizontal swarm postion (Figure 4C). For example, at the swarm peak, the horizontal velocities at the swarm surface and floor are twice the swarm speed and close to zero respectively (Figure 4C, Supplementary Figure 9D). As in the model, our data show that the difference in cell motion at the swarm surface and floor has a clear relationship with swarm height, but not with the distance from the swarm peak (Figure 4F), which has a consistent lag behind the bacterial peak (Figure 2B & 2C). In confirming model predictions, our results strongly suggest that the material properties of an active fluid – viscosity, friction, activity and surface tension – emerge in large numbers of migrating cells, resulting in emergent flow-fields that dictate cell organization.

The model predicts a reduction in cell flow at the swarm core – do cells stop moving entirely, or lose directionality? Both situations are equivalent in our continuum model. Therefore to address this, and to go beyond the deterministic continuum interpretation of our model, we analysed the variance in single cell motion across the swarm cross section (Figure 4G, Supplementary Figure 9C & 9D). The mean speed of individual cells remains relatively constant throughout the swarm, with the exception of the swarm rear which tends to zero (Figure 4H, Supplementary Figure 9E). However, cell motion is uniform and directional at the swarm surface and, conversely, highly variable and predominately misaligned with the gradient at the swarm core regions (Figure 4G). This misalignment perhaps suggests cells in the swarm core are impeded from moving forward. This transition between order and disorder is summarised by a steady reduction from large Péclet numbers at the swarm surface, indicative of strongly directed motion, to a Péclet number of around 1 at the swarm core, indicative of a balance between directed and random motion (Figure 4H, Supplementary Figure 9E). Overall, these results imply that the reduction in average cell movement at the swarm core is caused by a transition from advective-to diffusive-dominant motion.

### Implications of active fluid behaviour for development

Migration of groups of cells is a widespread feature of developmental and disease processes, from morphogenesis to wound healing and metastasis (*29*). Here we have shown that emergent fluid-like properties of cell groups -clump shedding and vortex motion - together have implications for how cells organise themselves in space, which determines the decision of cells to remain in the undifferentiated state or to enter the developmental programme. Differing from conventional models of cell state allocation based purely on positional information (*30*) or cell autonomous fate allocation (*31*), our data suggest a different mechanism (Figure 5). The outcome for a single cell (to differentiate or remain undifferentiated) will depend on its location within the vortex at the time of droplet shedding. We refer to this mechanism of cell fate allocation as “musical chairs” decision-making. This is based on analogy to the party game in which there are fewer chairs than children. When the music stops (droplet shedding), the children not finding chairs are eliminated from the game (differentiation) whilst the other children get to stay in the game (with the continued possibility of feeding). We propose that similar mechanisms will arise in other developmental and disease contexts involving migration of groups of physically interacting cells in response to signaling gradients.

**Figure 5.**
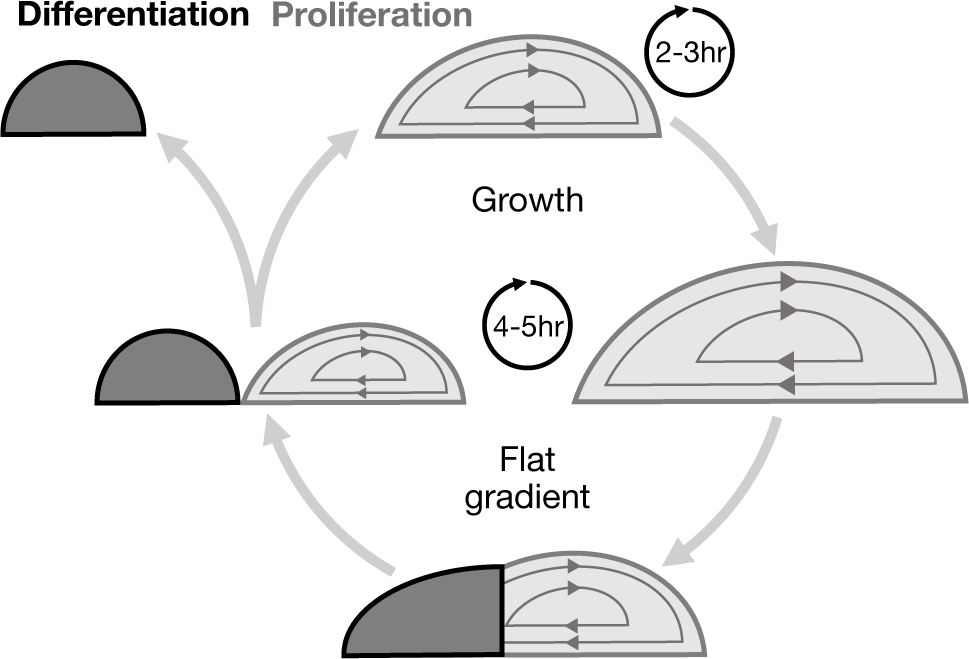
Musical chairs decision-making. Schematic of the properties of an active droplet that emerge during cell swarming, and influence cell differentiation and proliferation. Within the migrating swarm (light), cells proliferate and circulate with a maximum period of around 2-3hrs. Once a critical fraction of the swarm experiences a flat gradient (dark), then the cells at the swarm rear will collectively cease to migrate and deposit within a cell clump, which are destined for differentiation.

## Supporting information

Supplementary Methods (mathematical modelling)

## Acknowledgments

We are very grateful to Shu En Lim and Bobby Ford for hair donations, Adolfo Siairdi for providing the Dino-Lite, Olive Ford for help preparing feeding fronts, Ricciardo Barrientos, Chris Brimson, Ian Jones and Delan Alasaadi for discussions, Tim Rudder for useful interpretations of cell transport, and Philip Maini and Suraj Shankar for discussions about the mathematical model. Work was supported by Wellcome Discovery Award 226655/Z/22/Z to JRC. GLC was supported by an EPSRC Doctoral Prize Fellowship (EP/W524335/1). PP was supported by a UKRI Future Leaders Fellowship (MR/V022385/1). MPD was supported by the UK Engineering and Physical Sciences Research Council [EP/W032317/1]. BJW was supported by the Royal Commission for the Exhibition of 1851. Lightsheet imaging was supported by BBSRC (BB/R000441/1) to CJW.

## Index to Supplementary Information

1. **Supplementary Movies**
2. **Supplementary** Figures
3. **Methods**
4. **Mathematical Modelling**

**Video 1. Pattern formation during feeding front expansion (relates to Fig 1A)**

Macrophotography time lapse of wild type (top) and *acaA-* (bottom) *Dictyostelium* colonies. The feeding front appears as a ring that expands into the surrounging bacteria field. Cell clumps appear as circles that shed from the localised areas of swarm elongation. The *Dictyostelium* developmental programme organised by cAMP signalling is seen initially as a streaming pattern. Scale bars: 2mm.

**Video 2. Cell clump shedding dynamics (relates to Fig 1A’)**

Same as Video 1 but showing a different dataset obtained at a higher frame rate with a smaller field of view. Video shows the coursening of an irregular shaped cell clump into two circular clumps, that persist while surroudning isolated cells progress through the *Dictyostelium* developmental programme: formation of cell streams, tipped-mouns, migrating slugs and eventually fruiting bodies. Scale bars: 500µm.

**Video 3. Dynamics of cell clump shedding and gradient remodelling – top view (relates to Fig 1B)**

Maximum projection (birds’-eye-view) of light sheet imaging of feeding front dynamics showing penetration into the bacteria field and shedding of cell clumps. The cell nuclei are labelled in orange and the bacteria are labelled in green. Scale bar: 200µm.

**Video 4. Self-generated gradient dynamics – side view (relates to Fig 1C)** Maximum projection (top panel: side-view, bottom panel: birds’-eye-view) similar to Video 3 but with a different data set obtained at a higher frame rate with a. smaller field of view. The cell nuclei are labelled in orange and the bacteria are labelled in green. Scale bar: 100µm.

**Video 5. Quantification of swarm height and bacteria quantity (relates to Fig 2A)**

Quantification of the swarm height (top) and bacteria quantity (bottom) of the data shown in Video 3. The tick units are in <u>µm. See figure legend for</u> Fig 2A.

**Video 6. Model simulation**

Simulation of the active thin film model of directed swarm migration (Mathematical Supplement). Top: model predictions for swarm dynamics during shedding in the stationary reference frame of the lab. Colour map indicates the magnitude of the horizontal cell velocity. As in Figure 3B, black dots indicate respectively the front on rear of the swarm. Left: simulated swarm dynamics during shedding in the co-moving reference frame of the swarm front. Colour map indicates the magnitude of the horizontal cell velocity in the co-moving reference frame. Right: simulated swarm dynamics and chemoattractant profile (dark red line) during shedding in the co-moving reference frame of the swarm front. Also shown in dark red is the region where the gradient in the chemoattractant drops below a cell sensitivity threshold (Mathematical Supplement).

**Video 7. Periodic shedding of cell clumpes along lines of bacteria (relates to Fig 3C)**

Macrophotography time lapse of wild type *Dictyostelium* cells migrating along thin lines of bacteria, resulting in the periodic shedding of cell clumps (1 shedding event per 4.35h). Scale bar: 1mm.

**Video 8. High resolution imaging of the cell nuclei (relates to Figure 4)**

Cells at the top of the swarm move in more direction manner than cells at the floor. Same as Video 2, but showing a different dataset obtained at a higher frame rate and a smaller field of view for cell tracking, without imaging the bacteria. Scale bar: 100µm.

---

**Supplementary Figure 1.**
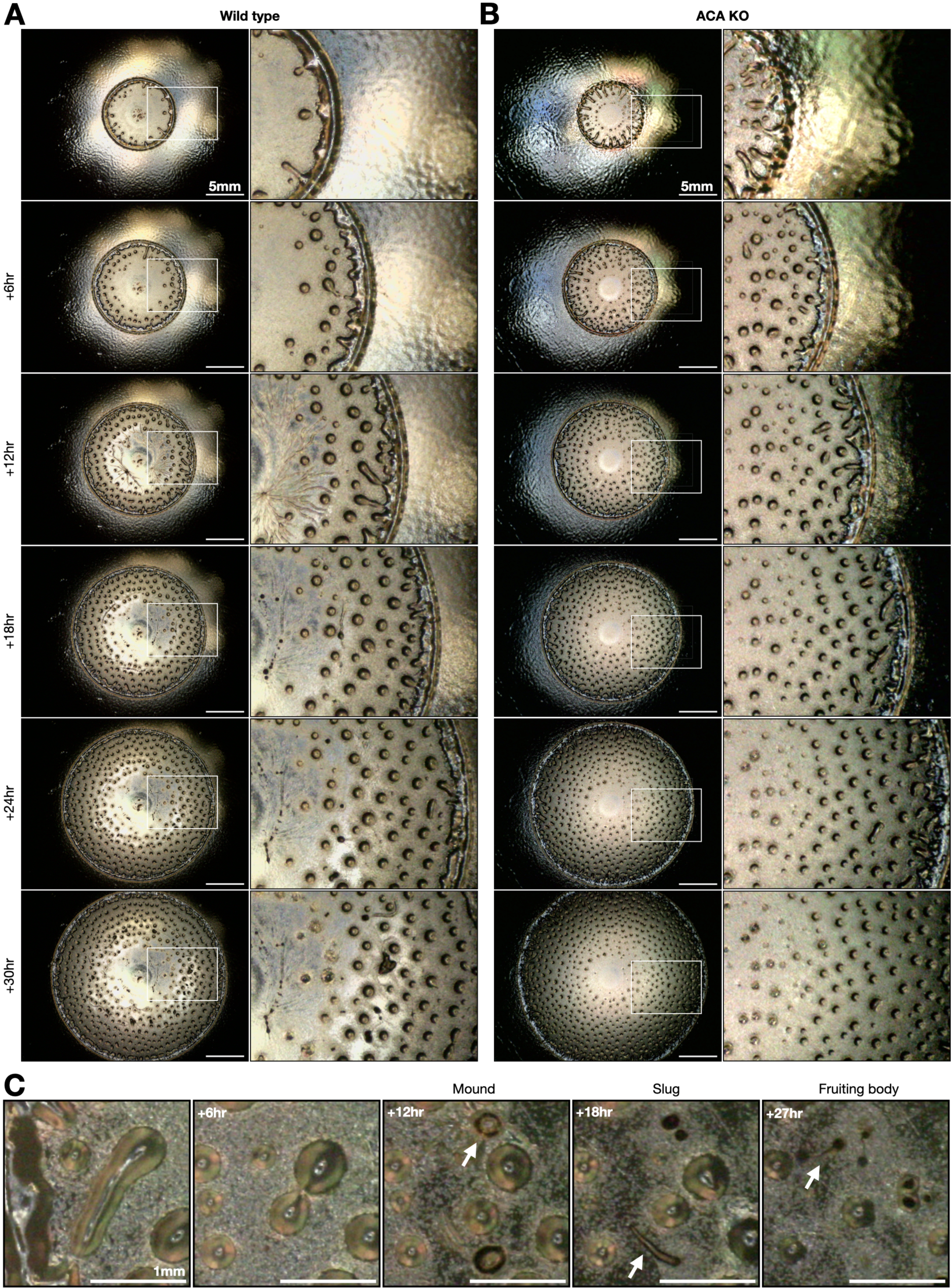
Patterns of cell clump shedding behind expanding feeding fronts. **A** Macrophotography of feeding fronts at 6h intervals, showing feeding front progression, patterns of clump deposition and late stage development. Scale bar 5 mm. **B** Feeding fronts of *acaA-* mutants showing similar feeding front progression and clump deposition but no later stage development. **C** Close up of wild-type cell clump progression, showing an irregular shaped clump pinching off the front then coarsening into two clumps. The clumps are persistent despite the occurrence of late stage development of surrounding cells.

**Supplementary Figure 2.**
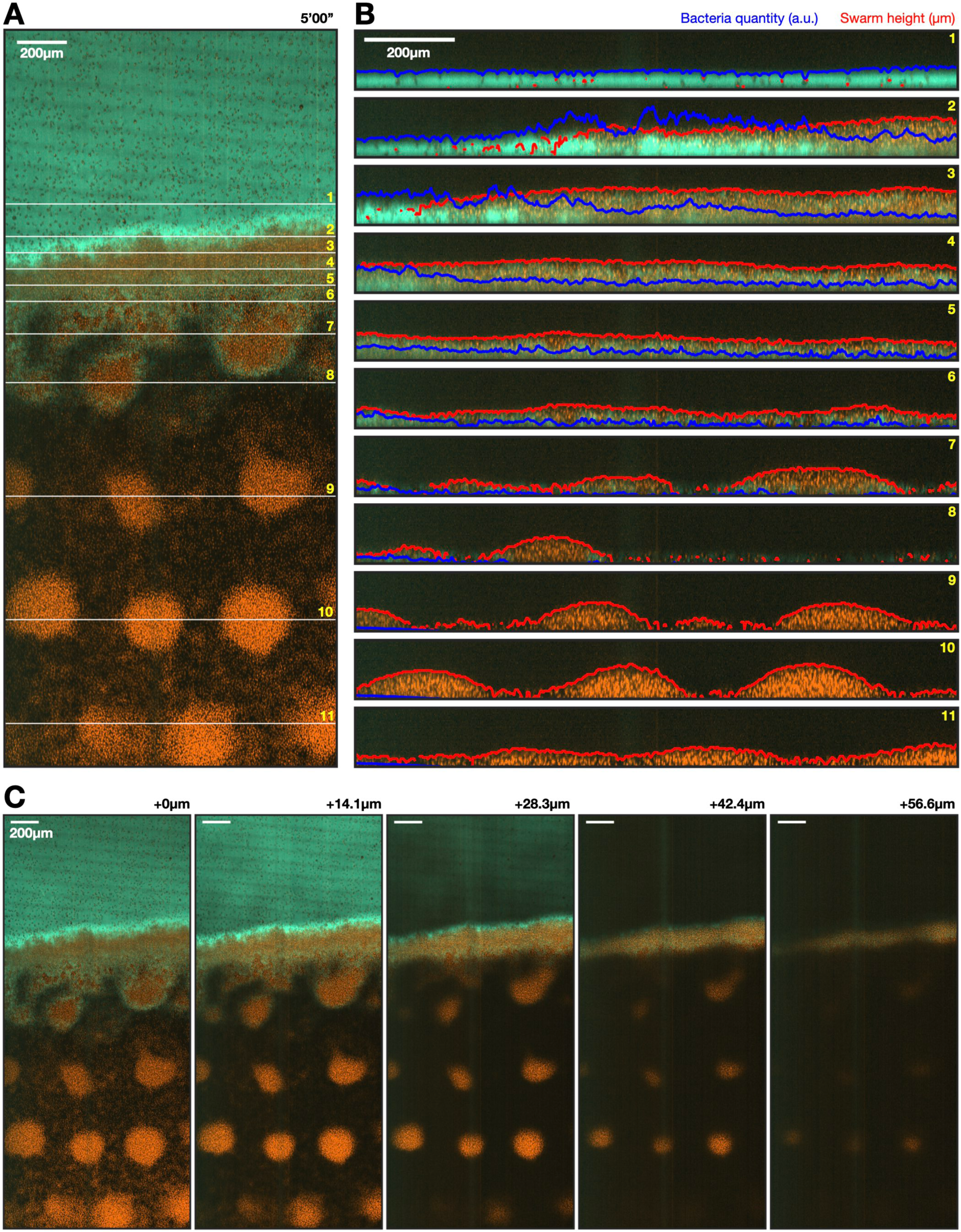
Quantification of swarm height and bacteria quantity from light sheet data. **A** Light-sheet imaging of *Dictyostelium* feeding fronts showing cell nuclei (orange) and bacteria (green). Shown is a single bird’s-eye-view of the entire imaging window **B** Side-view cross sections of the same data in A showing the measured swarm boundary (red line) and quantification of the bacteria (blue line). Each section corresponds to a line in panel A. **C** Same data as A but showing sectioning out of the plane.

**Supplementary Figure 3.**
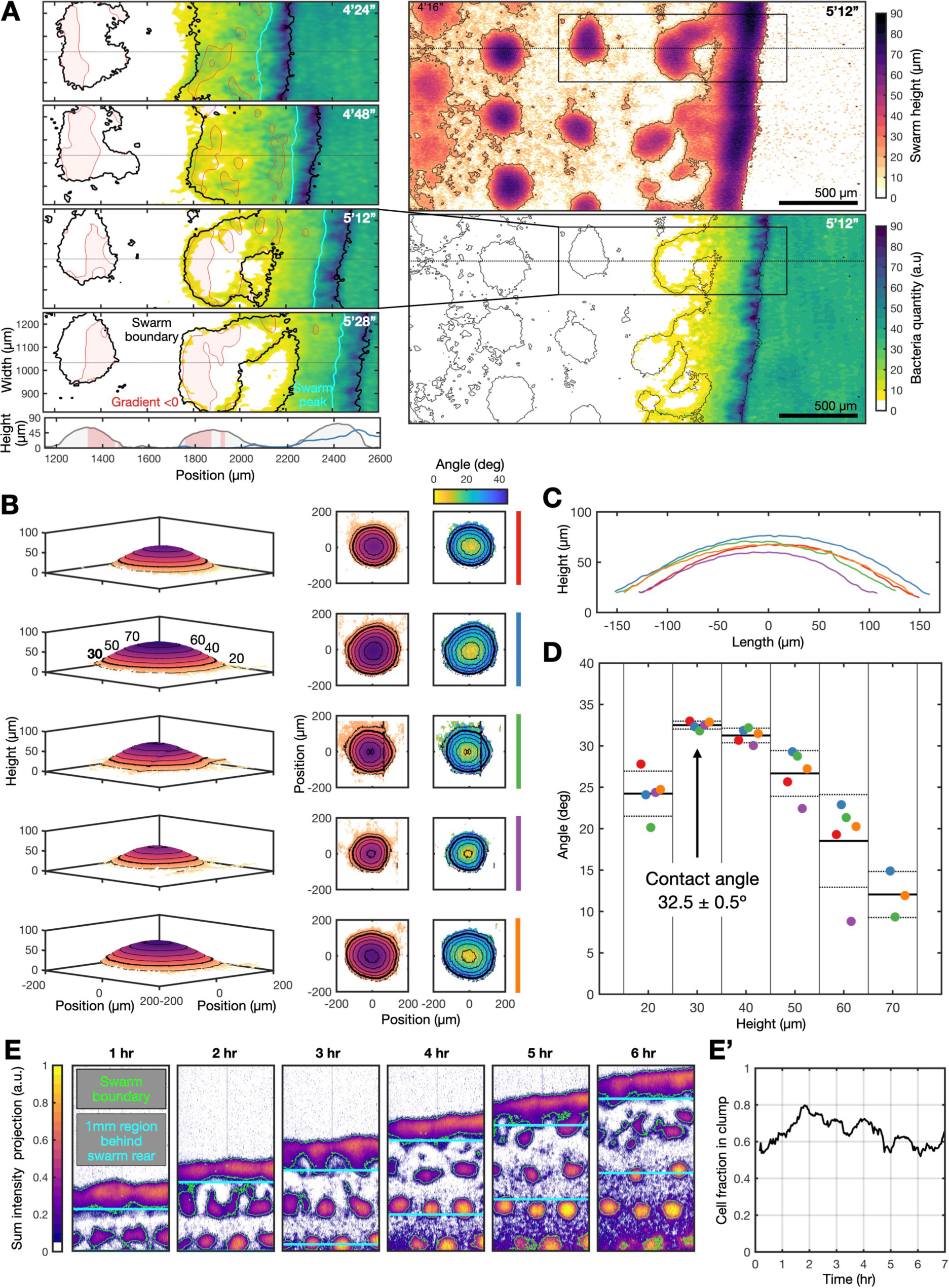
Quantitative features of cell clumps. **A** Quantification of the cell swarm height and bacteria quantity from the data shown in Fig 1B. The left column shows the distribution of bacteria throughout a shedding event, while highlighting the swarm boundary (where the height is 30µm) and peak and where the bacteria gradient is flattened (value less than zero). Right panels show the swarm height and bacterial quantity from the entire field of view. These data are heatmaps of the data in Figure 2A. **B** 3D plots, with contours, of the surface of five cell clumps (average over 60 min of imaging) from the data shown in Figure 1B. Also shown is the angle of the surface gradient (right column). **C** Cross sections of each clump in B, shown in different colours. **D** Scatter plots of the average angle of the surface gradient for each clump (colour), for different heights on the clump surface (x-axis). Also shown is the mean (thick line) and standard deviation (dotted line) for each height. The smallest height (30µm) at which the variance is negligible is highlighted and estimated as 32°-33°. **E** Proportions of cells in clumps and isolated cells. Sum intensity projections (cell nuclei) of the 3D images shown in Figure 1B and the fraction of cells that are left behind the feeding front within a clump as opposed to as single cells (**E’**). This value was quantified by comparing the total nuclei intensity within and outside clump boundaries (contours where the population height is 30µm, highlighted in green) in a 1mm region behind the swarm rear (highlighted in cyan).

**Supplementary Figure 4.**
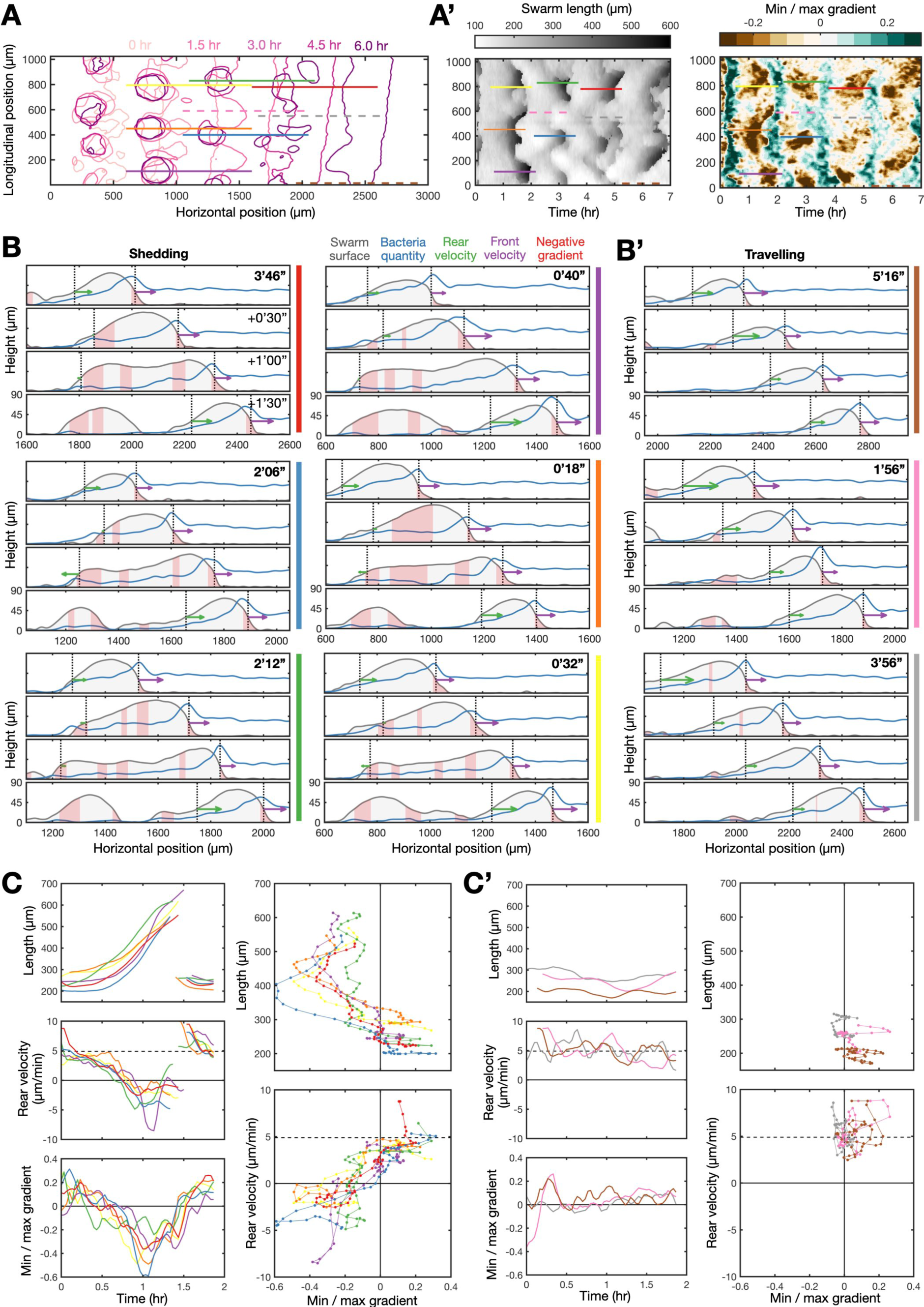
Swarm shape dynamics during travelling and shedding. **A** Contours of swarm and clump boundaries shown at 1.5 hr intervals (colour). **A’** Swarm length (left) and the ratio of the minimum and maximum gradient within the swarm (right) as a function of position across the imaging window and time. Coloured lines indicate the spatial and time points for the representative data show in in B. **B** Cross sections of a swarm over a 90 min period during the shedding phase (elongation and splitting) (**B**) and the travelling phase (no elongation) (**B’**). The colour bands refer to colours in A. These plots are the same representation as Fig. 2B, using a different sample point of the same dataset. **C** Elongation associated with a loss of positional information from the front due to a negative minimal gradient. Plots of the swarm length (top left), rear velocity (middle left) and the ratio of the minimum and maximum gradient (bottom) for the swarm cross sections shown in B (shedding phase). Also shown is the swarm length (top right) and rear velocity (bottom right) plotted against the ration of the mimum and maximum gradient. The dashed lines indicate the mean swarm speed. **C’** Same as C, but for cross section shown in B’ (travelling phase).

**Supplementary Figure 5.**
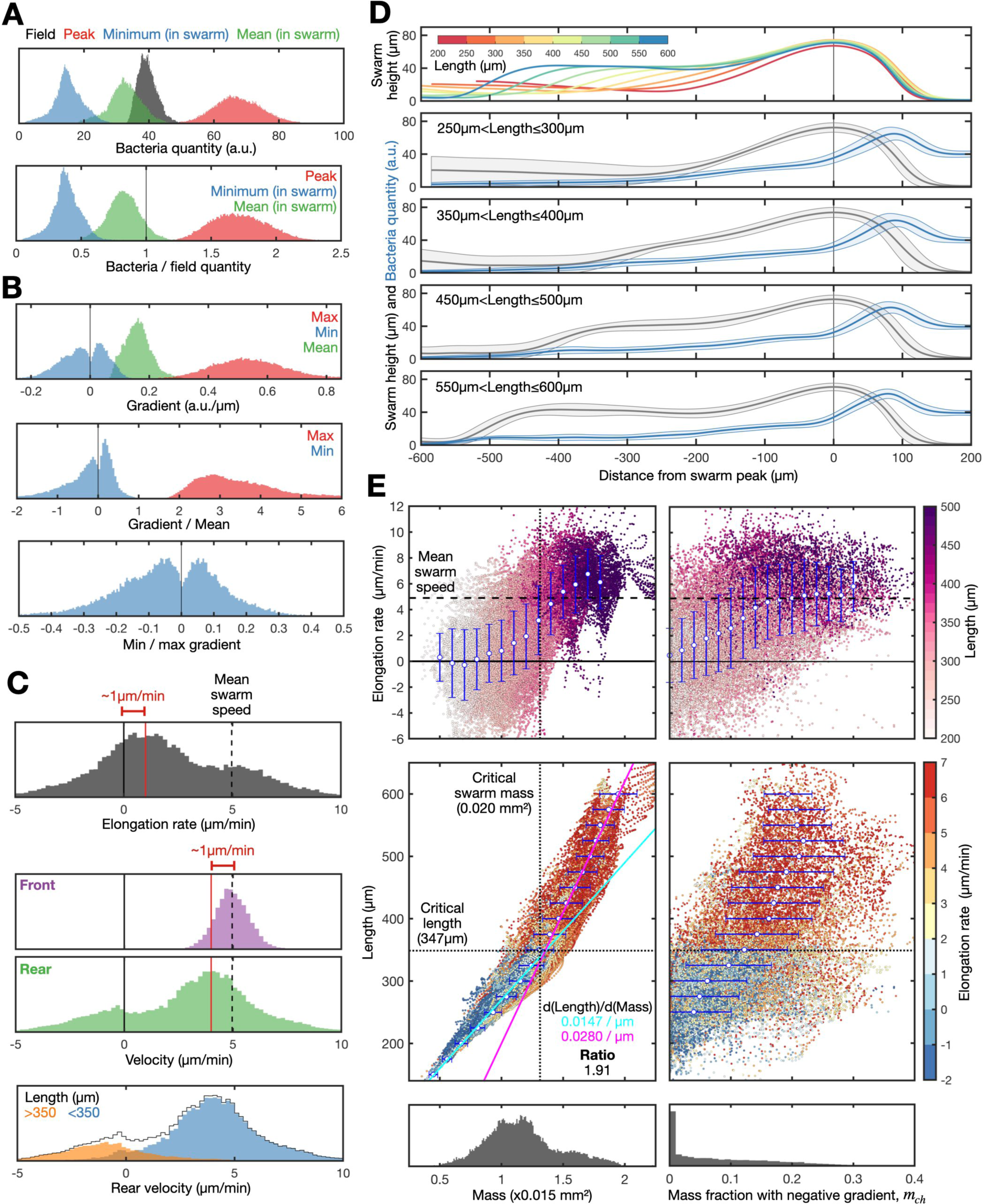
Extended analysis of cell swarm and signal gradient interactions. **A** Histograms of the minimum (blue) and mean (green) quantity of bacteria within the swarm boundary, and the quantity of bacteria at the bacteria peak (red) and bacteria field (black - averaged across 100µm-200µm ahead of the bacteria peak). Also shown are the minimum, mean and peak bacteria quantity relative to the quantity of the bacteria field (bottom). **B** Gradient statistics: histograms of maximum (red), minimum (blue, neglecting the minimum gradient at the front of the bacteria peak) and mean (green) value of the gradient within the swarm (top), the ratio of the maximum and minimum gradient to the mean gradient (middle), and the ratio of the minimum gradient and maximum gradient (bottom). The maximum value (at the front of the swarm) is consistently greater than twice the mean value of the gradient within the swarm. The minimum gradient is double-peaked around zero. Overall this indicates the gradient at the front is consistently positive whereas the gradient in the swarm fluctuates. **C** Shown are histograms of the elongation rate (black), showing two elongation states-slow and fast. Also shown are the front (purple) and rear (green) velocities, highlighting the mean swarm speed (dashed line) revealing the difference in the mean and rear swarm velocity-which accounts for the low baseline elongation. Bottom panel: histograms of the rear swarm velocity for regions where the swarm length is less than (blue) and greater (orange) than the critical swarm length (350µm) separating the “travelling” and “shedding” phases of swarm behaviour, shown in Figure 2F. **D** Low variance swarm and bacterial profiles at different swarm lengths, showing reproducible swarm dynamics. Top: mean swarm boundaries (taken from all spatial and time point) for swarms of different lengths (colour). Bottom: mean and standard deviation of the swarm boundaries (grey) and bacteria quantity (blue) for swarms of different lengths. **E** Comparing swarm dynamics, size and mass (area under the curve) with model predictions in Fig. 3F. Left panels show how elongation rate (top), swarm length (middle) relate to the mass (bottom). Below a critical mass, the length linearly increases (cyan line) with mass. Above this critical mass, the length (and elongation rate) increases at a faster rate (around twice as fast). Critical values of both mass (0.020 mm²) and length (347µm) were derived from the intersections between the linear fits. Right panels show the same analysis, comparing length and elongation with respect to the mass fraction of the swarm that experiences a negative gradient. A steady increase in the elongation rate (top) and length (middle) occurs as the mass fraction with a negative gradient increases.

**Supplementary Figure 6.**
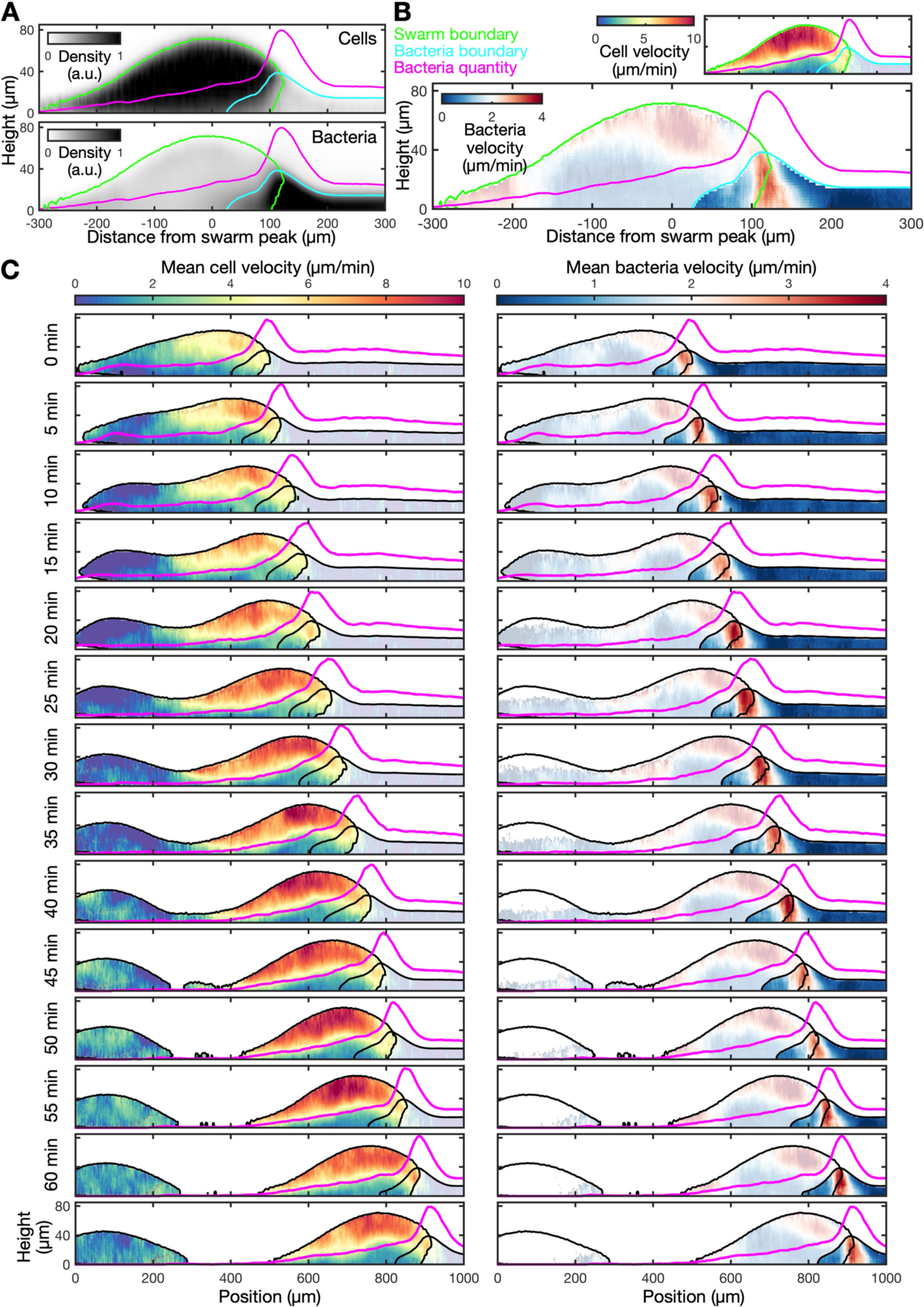
The “snowplough” model: particle image velocimetry of cell nuclei and bacteria. **A** Mean density of cells (top) and bacteria (bottom) across the face of the swarm (dataset shown in Fig 1C). Also shown are the boundaries of the swarm (green) and bacteria (blue), and the total sum of the bacteria along the swarm length (pink). **B** Velocity of bacteria is maximal at the leading edge of the swarm. Plots shows the velocity fields of bacteria and cells across the face of the swarm measured using PIV. **C** Velocity fields of bacteria and cells over a 1 h period during a shedding event.

**Supplementary Figure 7.**
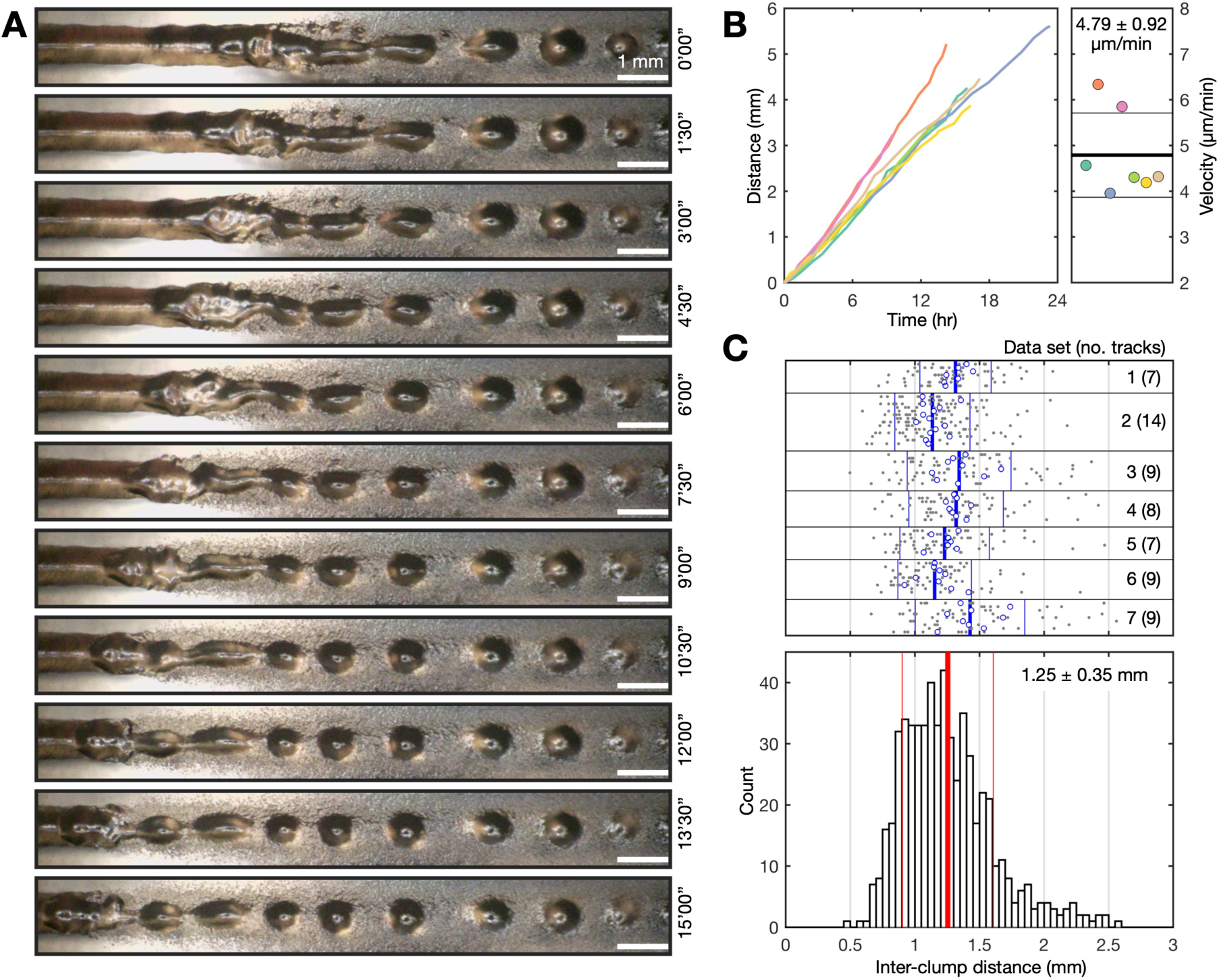
Estimating the periodic shedding rate of cell clumps. The shedding rate (1 clump per 4.35 hours) was estimated by dividing the swarm speed by the interclump distance of swarms on thin bacterial lines: **A** Time lapse macrophotography of an isolated and compact *Dictyostelium* swarm travelling along, and clearing, a thin line (∼500µm width) of bacteria. **B** Tracks of the position (left) and average speed (right) of *Dictyostelium* swarms travelling along bacteria lines (representative data shown in A). Seven biological repeats are shown in colour. **C** Quantification of the average distance between cell clumps generated by swarms travelling along bacteria lines. The experiments were performed by generating a set of bacteria lines on a single plate. 7 experiments are shown, 1 per box. Each grey dot within a box is the distance between clumps, with the mean (blue dots) of the inter-clump distance of invidiual lines, together with the mean (thick blue line) and standard deviation (thin blue line) for each experiment. The bottom panel shows a histogram of all measured inter-clump distances, together with the aggregated mean (thick red line) and standard deviation (thin red line).

**Supplementary Figure 8.**
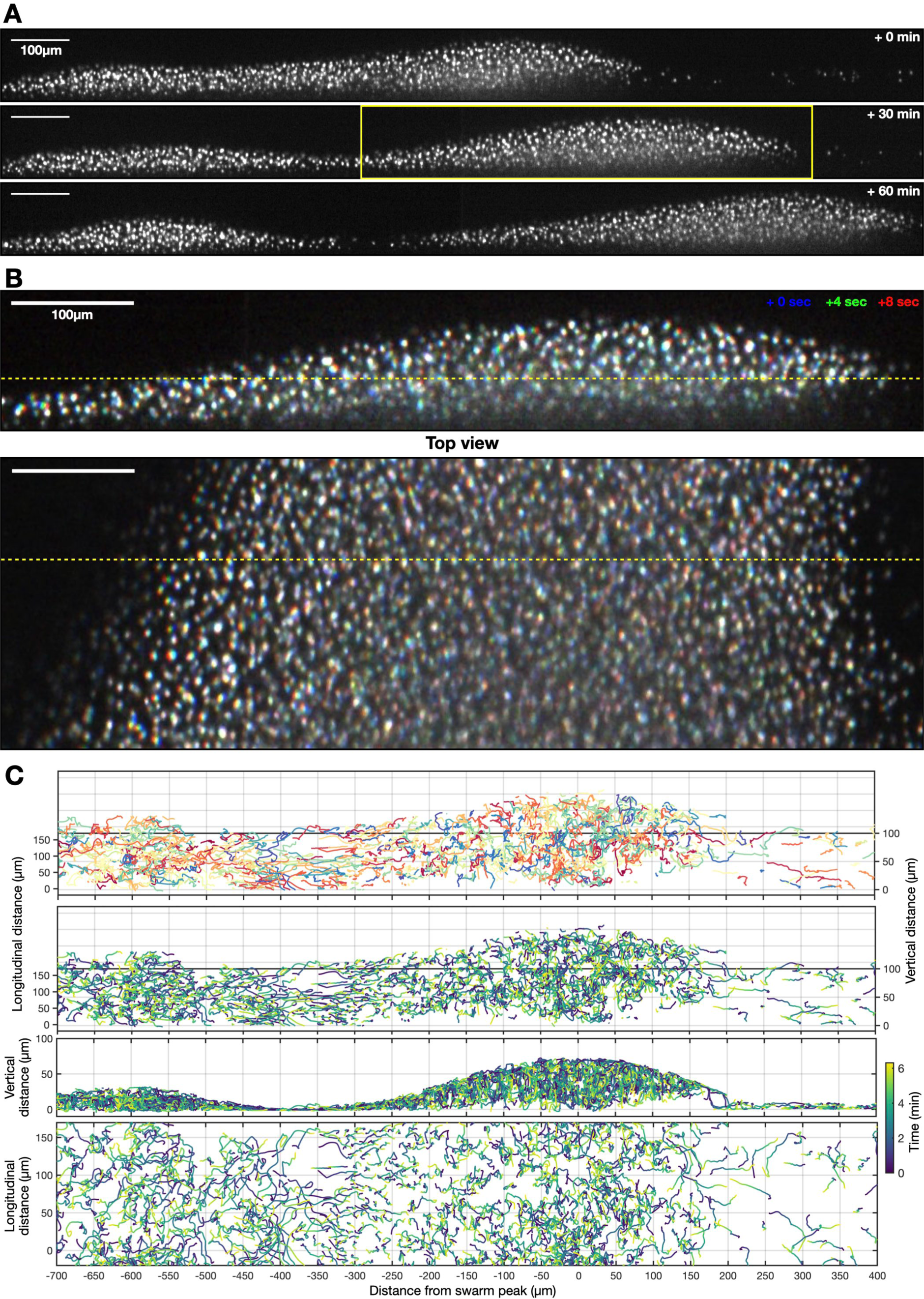
Tracking cell motion from high spatiotemporal resolution imaging of the swarm. **A** Cross section of the swarm (raw data) every 30 min during a shedding event. **B** Side and top views of the region of the swarm from the yellow box in A with three overlaid time points, captured at 4 s intervals. Different time points shown in different colours. The dashed yellow line in the top panel shows shows the vertical slice displayed in the bottom panel (and vice versa). **C** A sub-sample of 3D cell tracks coloured by their cell ID (top panel) and time (bottom three panels). Bottom three plots show (from top to bottom) the same 3D tracks from an alternative viewpoint (at a 45 degree angle) from the side and from the top.

**Supplermentary Figure 9.**
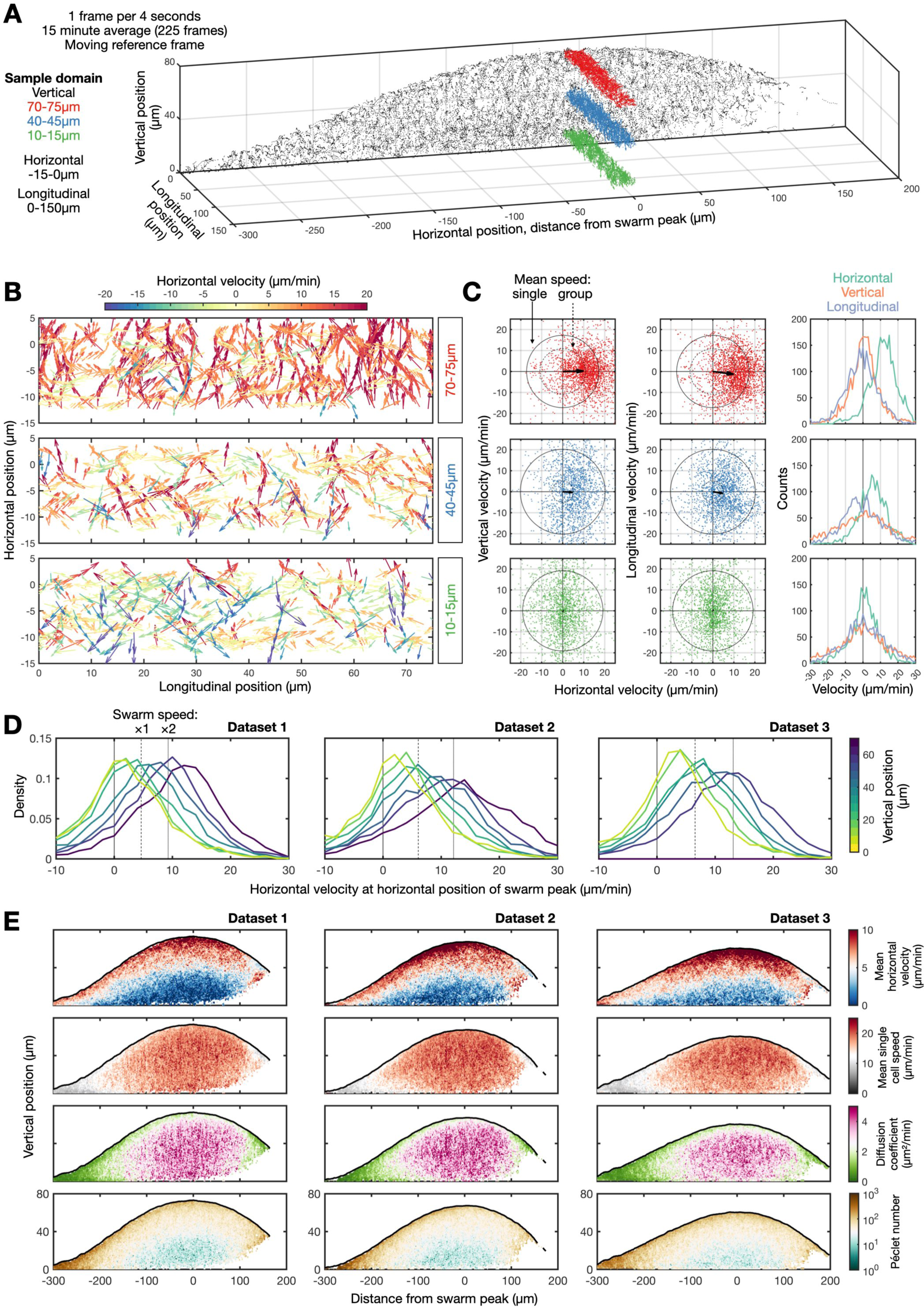
Quantification of cell motion within the swarm: finding the balance between advection and random motion. **A** 3D section of the swarm. In grey are individual cell coordinates from a thin slice of the swarm. The red, blue and green vectors show all measured velocities of cells at small longitudinal sections across the 3D volume. **B** Zooming in on the three coloured slices in A to show velocity vectors. C Scatter plots of the 3D velocities measured at the three different swarm heights. Black arrows indicate average motion. Circular lines are full or dotted. Full lines represent the average speed of single cells, which is relatively invariant. Dotted lines are the average speed of the group at that swarm position, which changed depending on swarm vertical position. Right panels show the distributions of the horizontal, vertical and longitudinal velocities at the three different locations. **D** Transition from directed to random motion at various vertical position in the swarms. These plots (from three biological repeats) show how the horizontal velocity (green curve in right panel of C) changes as a function vertical position in the swarm. At the top of the swarm, cells move at twice the swarm speed. At the bottom, their average horizontal velocity is close to zero. **E** Same representation as Fig. 4h for 3 biological repeats. Top panels show mean horizontal velocity across the swarm face. 2^nd^ row shows mean single cell speed. 3^rd^ row shows diffusion coeffcient and 4^th^ row shows the Péclet number.

## Methods

### Cell handling

We used *Dictyostelium* AX2 cells with red fluorescent nuclei generated by insertion of a histone H2B-mCherry gene into the *act5* gene (*32*). For routine culturing, cells were innoculated on lawns of *Klebsiella* on SM agar (*33*). To prepare feeding fronts for imaging, 120µl of a *Klebsiella* suspension was evenly spread across a 9 cm agar plate containing dilued SM (1 SM: 9 KK2; 1.5% agar) left to almost completely dry before seeding a *Dictyostelium* colony. Care was taken to generate an even and smooth bacteria lawn. For fluorescent imaging of bacteria, we used GFP-labelled Klebsiella (*34*). *Dictyostelium* colonies were seeded by resuspending around 10^7^ cells from the leading edge of an initial SM colony into 0.1 mL of KK2 buffer (20mM KPO_4,_ pH 6.0). 1 µl of this suspension was spotted onto the centre of the bacteria lawn. To prepare thin lines of bacteria, we used a human hair (made hydrophilic by cleaning in household detergent for 10 minutes and 12% NaClO for 20 min, with an ethanol wash between each use of the hair) dipped into bacteria culture, tapped dry, and then gently pressed against an agar plate. *Dictyostelium* cells was then spotted at the base of the printed lines of bacteria. For generating the *acaA* mutant cell lines, we replaced the hygromycin selection cassette in a published *acaA* targeting vector, pPPI725 (*11*) with a blasticidin resistance cassette from pDM1079 (*32*) by swapping NheI/NotI fragments. The targeting vector was linearised for transformation with NgoMIV. Transformation, selection and screening were carried out as described (*35*).

### Live cell imaging

For macrophotography (Fig 1A, Supplementary Figure 1 & 7, Video 1 & 2), a Dino-Lite USB microscope was used to image feeding fronts of *Dictyostelium* (1-2 days after innoculation) at 22°C (*16*). The sample was imaged every 2 minutes for 2-3 days, illuminating the sample only during image acquisition. To prevent desiccation, samples were imaged in a custom-built humid chamber – a completely dark and enclosed box, except for a hole at the top for imaging, with a platform (sample mounting) surrounded by a water reservoir. Macrophotography imaging data was analysed manually.

To 3D live image both bacteria and *Dictyostelium* cells across feeding fronts of *Dictyostelium*, we used a 3i Marianas light-sheet microscope (Dual Inverted Selective Plane Illumination Microscope, diSPIM) (*36*). Illumination and imaging were carried out above the sample at 45° to the surface with oil-dipping 10 x objectives. Samples were submerged in silicone oil. Imaging data were collected at 3 different spatiotemporal scales. Data in Figure 1B & 2, Supplementary Figure 2-5 and Video 3 & 5 were obtained by moving the sample 3mm through the light sheet along the axis parallel to feeding front travel at 2 µm step sizes every 2 minutes, imaging with both red (nuclei) and green (bacteria) light. The total volume of the field-of-view was 3000µm x 1300µm x 200µm with voxel dimensions 1.3µm x 1.3µm x 2 µm (width x length x height). Data in Figure 4, Supplementary Figure 8 & 9 and Video 8 were obtained by moving the sample 150 µm downwards through the light sheet at 1 µm step sizes every 4 seconds, imaging just the red nuclei. The data presented in Figure 1C, Supplementary Figure 6 and Video 4 were obtained by moving the sample 500 µm through the light sheet, perpendicular to the direction of swarm travel, at 1 µm step sizes every 15 seconds, imaging both green and red channels. The total volume of the field-of-view was 150µm x 1300µm x 200µm with voxel dimensions 1.3µm x 1.3µm x 2 µm (width x length x height). Slidebook2022 was used to deskew the imaging data and export to tiff format for downstream analysis.

### Image analysis

To quantify swarm shape, the upper and lower surfaces of both the *Dictyostelium* and bacteria populations were calculated using Matlab’s edge detection algorithm applied to binarised images of the cross sectional plane perpendicular and parallel to the direction of travel. The surface of the agar was determined at each time point by fitting a plane to the bottom surfaces of the bacteria and *Dictyostelium* populations. The quantity of bacteria was estimated by a sum z-projection. The location of the swarm front and the rear were defined as the positions where the swarm height was 30µm. The bacteria gradient was estimated by the spatial derivative of the total amount of bacteria across a distance of 6 cell widths (6 x 13µm). The mean and minimum values of the bacteria gradient were calculated as the mean and minimum values between swarm peak and rear.

To estimate the flow field of bacteria (Supplementary Figure 6), particle image velocimetry (using PIVlab, Matlab) was applied to the bacteria (green) and cell nuclei (red) channels of each 2D plane (parallel to the direction of swarm travel) and then averaged (500µm) at each time point.

To estimate cell flow fields within the swarm, individual nuclei were first identified by watershed segmentation (SCF-MPI-CBG Fiji update site) of Gaussian and then median filtered (3D) raw images. The centroid of each labelled nuclei was used for cell tracking (TrackMate: simple LAP tracker, CSVImporter). Cell velocities were determined by the second order central finite difference of cell positions. The mean cell velocity field across the swarm was calculated by averaging the velocity of each cell relative to the peak of the swarm (4µm (length) x 2µm (height) grid), averaged over a 15 minute period.The mean cell velocity field in the moving reference frame of the swarm was determined by subtracting the swarm velocity from the mean cell velocity field. The streamlines were determined by the Matlab streamline function. The Péclet number *P*_2_ = *L u*/ *D* was calculated at each point in the travelling reference frame. Variable *u* (µm/min) is the mean cell speed (3D). Variable *L* = *A*/*S* (µm) is the characteristic length, defined as the area of the swarm (viewed from the side), *A*, divided by the length of the curve that defines the swarm surface, *S*. Variable *D* = σ^9^ *Δt*/6 (µm^2/min) is the diffusion coefficient at each grid point, where σ^9^is the variance of the instantaneous cell velocities (mean of the squared speeds relative to mean speed) at the grid point and *Δt* = 4 sec is the time between frames.

## Notes

### Competing Interest Statement

The authors have declared no competing interest.

