## Supplementary Methods (mathematical modelling) for "Pattern formation along signaling gradients driven by active droplet behaviour of cell groups"

### Supplementary material for “Pattern formation along signaling gradients driven by active droplet behaviour of cell groups”

Hugh Z Ford[X1,2], Giulia L Celora[X1,3], Elizabeth R Westbrook[1,2],
Mohit P Dalwadi[1,3], Benjamin J Walker[1,3,4], Hella Baughmann[5], Cornelis J. Weijer[6],
Philip Pearce[*1,3] & Jonathan R Chubb[*1,2]

[1]Institute for the Physics of Living Systems, University College London, UK
[2] Laboratory for Molecular Cell Biology, University College London, UK
[3] Department of Mathematics, University College London, UK
[4] Department of Mathematics, University of Bath, UK
[5] Intelligent Imaging Innovations Ltd, London, UK
[6] School of Life Sciences, University of Dundee, UK

[*]Corresponding authors
[X]Co-first authors

### 1 Mathematical Model

We derive a continuum mathematical model for the migration of Dictyostelium groups based on the hypothesis that physical cell-cell interactions via the formation, attachment and retraction of pseudopods result in cell groups having the following fluid-like material properties:

1) surface tension, $\kappa$, resulting from cell-cell adhesions [11];

2) viscosity, $\eta$, resulting from the transience of cell-cell adhesions [2];

3) bulk activity, $\xi$, generated by cells actively contracting as they move within the group [11, 7].

Here, we develop a minimal mathematical framework to model living droplets with a constant surface tension and viscosity, and a bulk activity that depends on a self-generated signal gradient. We aim to understand how these material properties contribute to the propagation and shedding of Dictyostelium groups during their directed migration in response to self-generated gradients in bacterial concentration.

#### 1.1 Full dynamical model of cell group migration: Pseudo-2D active thin-film model

We consider a $(x, y)$ cross-section of the group and describe it as a thin 2D active polar fluid film (see Figure SM1). As shown in Figure SM1, the Dictyostelium group is described by the location of its free surface, *i.e.*, its height $y = h(x, t)$, with units of microns. The velocity field $\boldsymbol{u}_c$, with units of microns/minutes, describes the internal flow of Dictyostelium cells within the film. This represents a locally averaged velocity of Dictyostelium cells at each point in space and therefore captures the strength of any directed motion, which we assume to be driven by surface capillary forces (related to the effective surface tension $\kappa$) and bulk active forces that are regulated by the chemotactic signal, *i.e.*, the magnitude of chemoattractant gradients.

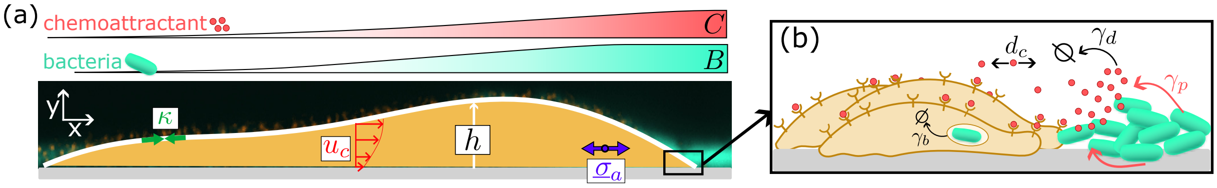


Figure SM1: Schematics illustrating the model set-up and the mechanisms included. (a) The Dictyostelium group is represented as an active thin film of height $h(x,t)$. The flow of cells in the group is described by the flow field, $\boldsymbol{u}_c$, which is driven by an emergent surface tension ($\kappa$) and an emergent bulk contractile active stress ($\underline{\sigma}_a = \xi S(\boldsymbol{e}_x \otimes \boldsymbol{e}_x)$). Similar to classic lubrication theory, the velocity field is dominated by its horizontal component, which follows a Poiseuille-like profile (see Section 1.1.2). The scalar fields $B$ and $C$ indicate, respectively, the local concentration of the bacteria cells and the chemoattractant molecules produced by the bacteria (*e.g.*, folic acid). (b) Dictyostelium cells move up self-generated gradients in the concentration of chemoattractant molecules (red particles) that are produced by the bacterial cells. As Dictyostelium cells migrate, they feed on and deplete the bacterial population, thereby shaping chemoattractant gradients.

Because the group height ($\sim 10\,\mu$m) is significantly smaller than its characteristic length ($\sim 100\,\mu$m), we adopt a lubrication approximation [9, 1, 14]. For brevity, we present the reduced form of the governing equations obtained at the leading order in the lubrication limit since the derivation of the model via asymptotic methods follows standard approaches [9, 14].

##### 1.1.1 Bacteria and chemoattractant concentration dynamics

We model the overall amount of bacterial cells at a given spatial location $x$ via a continuous concentration field $B = B(x,t)$, with units cells/microns, which is consumed by cell groups at a rate proportional to its height (see Figure SM1b). Neglecting for simplicity any effect due to the advection of bacteria by the cell flow within the group, $B$ evolves according to the following spatially-structured ODE:

$$\frac{\partial B}{\partial t} = -\gamma_b B h. \tag{1}$$

In Eq. (1) the term $\gamma_b h$, with units of 1/hour, is the rate of bacterial consumption within the cell group, where $\gamma_b > 0$ is a constant and $h(x,t)$ is the height of the cell group, as defined previously.

The chemoattractant concentration $C = C(x,t)$, with units of nanomolars, is modelled as a diffusible species with diffusion coefficient $d_c$. It decays at a constant rate, $\gamma_d$, while being produced at a rate $\gamma_r$ by bacteria cells (see Figure SM1b). We assume the chemoattractant diffuses rapidly in the $y$-direction, in line with expected diffusive timescales in the lubrication limit. We neglect any advection of the chemoattractant by Dictyostelium cell movement. The time evolution of $C$ is dictated by the following reaction-diffusion partial differential equation:

$$\frac{\partial C}{\partial t} = d_c \frac{\partial^2 C}{\partial x^2} - \gamma_d C + \gamma_p B. \tag{2}$$

The direction of droplet movement is dependent on the initial positioning of the bacterial lawn, *i.e.*, the initial condition for $B$, which introduces some pre-patterning that breaks the left-right symmetry in the system. We discuss this further in Section 1.2.

##### 1.1.2 Dictyostelium cell movement

We assume that the total stress within the cell group

$$\underline{\sigma} = -p\underline{\mathrm{I}} + \eta[\nabla \boldsymbol{u}_c + (\nabla \boldsymbol{u})^T] + \xi S\left(\boldsymbol{e}_x \otimes \boldsymbol{e}_x - \frac{1}{2}\underline{\mathrm{I}}\right) \tag{3}$$

is the sum of three contributions: the pressure $p$, a viscous stress with shear viscosity $\eta$, in units of Pascals·minutes, and an active stress with constant activity $\xi > 0$, in units of Pascals. In defining the active component of the stress (final bracketed term on the right-hand side of Eq. (3)), we have assumed any nematic order (or cell polarity) to be directed along the vector $\boldsymbol{e}_x$, *i.e.* in the direction of the chemoattract gradient. The strength of the nematic order parameter $S \in [0, 1]$ depends on the magnitude of the chemoattractant gradient and is discussed in detail below. In the lubrication limit, the stress in the fluid is dominated by the pressure $p$, which is obtained by imposing force balance at the air-cell group interface along the direction normal to the free-surface [1]:

$$p = -\kappa \left[ \frac{\partial^2 h}{\partial x^2} - \Psi(h) \right] - \frac{\xi S}{2}. \tag{4}$$

The first two terms on the right-hand side of Eq. (4) are common in the modelling of passive droplets and capture respectively the capillary forces, which result from an effective constant surface tension $\kappa$, in units of Netwons per microns, and a disjoining pressure resulting from the interaction of cells with the surface, which gives rise to an emergent macroscopic contact angle ($\theta_e$). We here use a standard form for $\Psi$ [1]:

$$\Psi(h) = \frac{3h_\delta^2 \tan^2 \theta_e}{(h + h_\delta)^3} \left( 1 - \frac{h_\delta}{h + h_\delta} \right). \tag{5}$$

In Eq. (5) $\theta_e$ is the equilibrium contact angle for the film and $h_\delta$ is a correction introduced to allow for dewetting without introducing a pre-wetting layer. Here, we take $h_\delta$ to be half the size of a Dictyostelium cell ($h_\delta = 5\mu m$) so that $\Psi$ describes effective attractive interactions between the layer of cells in contact with the floor and the floor itself. These forces allow for film rupture when cell layers are extremely thin [10] ($h < h_\delta$), but otherwise have a negligible effect. For a static film, Eq.(5) fixes the contact angle with floor to be $\theta_e$; however, for a moving thin film, Eq.(5) dynamically regulates the contact angle.

We hypothesise that cell-floor interactions allow cells to move along the floor while experiencing effective friction; we model these effects by imposing a standard Navier slip condition at the floor [9]:

$$u_c - \ell_s \frac{\partial u_c}{\partial y} = 0, \quad y = 0, \tag{6}$$

where $\ell_s > 0$, with units of microns, represents the slip length, which is inversely proportional to the cell-floor friction. With these assumptions, we find that $u_c$ follows a leading-order parabolic profile:

$$u_c(x, y, t) = \frac{F(x,t)}{\eta} \left[ h(x,t)\,(\ell_s + y) - \frac{y^2}{2} \right]. \tag{7a}$$

The function $F$ describes the net force that drives cell movement; this consists of three terms:

$$F(x,t) = \kappa \frac{\partial^3 h}{\partial x^3} - \kappa \frac{\partial \Psi(h)}{\partial x} + \xi \frac{\partial S}{\partial x}. \tag{7b}$$

The profile of the horizontal velocity in Eq. (7) is obtained by integrating the $x$-component of the momentum conservation equation ($\eta \partial_{yy} u_c = \partial_x \left[ p - S\xi/2 \right]$), balancing surface forces in the direction tangential to the air-cell group free-surface ($\partial_y u_c|_{y=h(x,t)} = 0$), and imposing the slip condition (6) within the lubrication limit.

For a passive droplet to move, an external force field (such as gravity) is needed. By contrast, active droplets can self-propel because energy is injected into the system at the microscale. In our specific system, active forces are generated through cell-cell interactions at the microscopic level, which result in an effective active stress at the population level. The constant $\xi$ in Eq. (7b) is the activity parameter that measures the typical size of the active stresses. Generally, $\xi$ can be of either sign. Using the same sign convention as in [14], here we take $\xi > 0$, under the assumption that Dictyostelium cell activity generates contractile forces in the preferred direction of motion through pseudopods that temporarily attach to other neighbouring cells [7, 3].

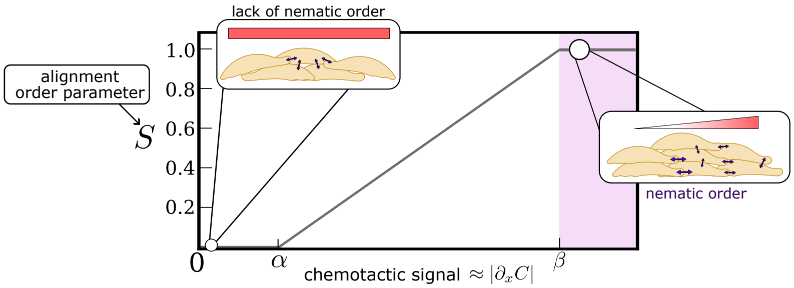


Figure SM2: Modulation of the alignment function $S$ in Eq. (7b) by chemoattractant gradients. If chemoattractant gradients are below the sensitivity threshold $\alpha$, cells lack information regarding the location of the bacteria and their movement is random ($S = 0$). When chemoattractant gradients are in the range $[\alpha, \beta]$, the bias of cell movement is in the direction of the chemoattractant gradients and the alignment function increases linearly until it saturates at $S \approx 1$, which corresponds to perfect bias of cell movement in the direction of increasing bacteria concentration.

Practically, Dictyostelium cells achieve directed migration by biasing the distribution of pseudopods [7]. In the absence of any directional information, the distribution of pseudopods, and therefore cell movement, is random. Chemotaxis is thought to bias random cell motility, by favouring retention of the pseudopod that experiences the higher attractant concentration. The stronger the chemotactic signal, the stronger the bias. In our model, chemotaxis bias is introduced by coupling the order parameter $S \in [0, 1]$ to the chemotactic signal. When $S = 0$, cell movement is random and the collective behaves effectively as a passive fluid. When $S = 1$, cell movement is strongly biased and the strong alignment in the direction of cell contractility results in maximal active stresses [12]. For simplicity, we assume a linear dependence of the local alignment parameter on the chemotactic signal, here generally denoted by $\omega$. To account for the loss of positional information when the response function is small, and for the saturation on the local strength of alignment when all cells are completely polarised, we assume that the alignment strength is a saturating ramp function of its input, $\omega$ (see Figure SM2)

$$S_{\alpha,\beta}(\omega) = \begin{cases} 0, & \omega < \alpha, \\ \frac{\omega-\alpha}{\beta-\alpha}, & \alpha \leq \omega < \beta, \\ 1, & \omega \geq \beta, \end{cases} \tag{8}$$

where $\beta > \alpha > 0$ are constants that control the width of the interval in which the order parameter $S$ increases linearly and its gradient. In practice, Dictyostelium cells will sense gradients in receptor occupancy, which can yield random cell movement at high chemoattractant concentrations as a result of receptor saturations [15, 4]. Here we assume that the chemoattractant concentration remains below the saturation threshold of the binding sites ($\approx 20\,\text{nM}$ [15]), so that cells respond to gradients in the chemoattractant, and set $\omega = |\partial_x C|$ (see Figure SM2). Since the model uses higher order derivatives of $S$, we adopt a smoothed approximation of (8) in practice (see Section 1.2).

###### 1.1.3 Dictyostelium group dynamics

The time-evolution of the height function $h$ is determined by imposing mass balance. Given that the density of cells within the droplet is assumed to be constant, we find that:

$$\frac{\partial h}{\partial t} + \frac{\partial Q}{\partial x} = r\frac{B^2}{m_B^2 + B^2}h, \tag{9a}$$

where the flow rate $Q$ is obtained by integrating the horizontal cell velocity in Eq. (7a) over $y$:

$$Q(x,t) = \int_0^h u_c(x,y,t)\,dy = \frac{F(x,t)}{\eta} h^2(x,t) \left[ \ell_s + \frac{h(x,t)}{3} \right]. \tag{9b}$$

In Eq. (9a), the rate of cell proliferation is taken to be a non-linear function of the concentration of bacteria. Cells proliferate at their maximum rate ($r > 0$, with units 1/minutes) as long as there is enough food, while they arrest when starved. Here, $m_B > 0$, in units of cells/microns, indicates the concentration of bacteria below which cell proliferation is arrested.

###### 1.1.4 Non-dimensional model

We non-dimensionalise the governing equations using the following scalings:

$$\begin{aligned} x = L\hat{x}, \quad y = \tan\theta_e L\hat{y}, \quad t = \frac{L}{U}\hat{t} \quad u_c = U\hat{u}_c, \\ h = \tan\theta_e L\hat{h}, \quad B = B_\infty \hat{B}, \quad C = \frac{\gamma_p B_\infty}{\gamma_d}\hat{C}, \end{aligned} \tag{10}$$

where $L$ is the characteristic length of the cell group, $B_\infty$ is the characteristic concentration of bacteria, $U$ is the characteristic velocity of the cells within the cell group and $\theta_e$ is the equilibrium contact angle. By substituting Eqs. (10) into Eqs. (1)-(2) and (7)-(9), we obtain the following non-dimensional system of coupled partial differential equations for $\hat{B}$, $\hat{C}$ and $\hat{h}$:

$$\textbf{Bacteria consumption} \;:\; \frac{\partial \hat{B}}{\partial \hat{t}} = -E\hat{B}\hat{h}, \tag{11a}$$

$$\textbf{Diffusion and production of chemoattractant} \;:\; \frac{\partial \hat{C}}{\partial \hat{t}} = D_c \frac{\partial^2 \hat{C}}{\partial \hat{x}^2} - \Gamma_d \left( \hat{C} - \hat{B} \right), \tag{11b}$$

$$\textbf{Dynamics of the thin-film height} \;:\; \frac{\partial \hat{h}}{\partial \hat{t}} = -\frac{\partial}{\partial \hat{x}} \left( M(\hat{h}) \frac{\partial \pi}{\partial \hat{x}} \right) + \frac{R\hat{B}^2}{\hat{m}_B^2 + \hat{B}^2}\hat{h}, \tag{11c}$$

where the mobility $M$ and the pressure $\pi$ are defined as:

$$M(\hat{h}) = \frac{1}{Ca_\kappa} \hat{h}^2 \left( \frac{\hat{h}}{3} + L_{sd} \right), \tag{11d}$$

$$\pi = \frac{\partial^2 \hat{h}}{\partial \hat{x}^2} - \frac{3H_\delta^2}{(\hat{h} + H_\delta)^3} \left( 1 - \frac{H_\delta}{\hat{h} + H_\delta} \right) + Ca_\xi S_{\hat{\alpha},\hat{\beta}} \left( \frac{\partial \hat{C}}{\partial \hat{x}} \right), \tag{11e}$$

In Eq. (11e), the pressure accounts for three contributions: 1) the capillary pressure, 2) the disjoining pressure and 3) the active pressure, which is mediated by the order parameter $S$, defined in Eq. (8), with rescaled hyperparameters $\hat{\alpha}$ and $\hat{\beta}$. In the non-dimensional form of the model, the evolution of the cell group height $\hat{h}$ is determined by seven non-dimensional parameters:

$$\begin{aligned} L_{sd} = \frac{\ell_{sd}}{H}, \quad Ca_\kappa = \frac{U\eta L^3}{\kappa H^3}, \quad Ca_\xi = \frac{\xi L \gamma_p B_\infty \bar{S}'}{2\gamma_d H \kappa}, \quad R = \frac{rL}{U}, \\ H_\delta = \frac{h_\delta}{L\tan\theta_e}, \quad \hat{\alpha} = \frac{\alpha L \gamma_d}{\gamma_p B_\infty}, \quad \hat{\beta} = \frac{\beta L \gamma_d}{\gamma_p B_\infty}, \end{aligned} \tag{12a}$$

where $\bar{S}' = \max_{\omega \in [0,\infty)} S'(\omega)$ and three non-dimensional parameters determine the evolution of $\hat{B}$ and $\hat{C}$:

$$D_c = \frac{d_c}{UL}, \quad E = \frac{eLH}{U}, \quad \Gamma_d = \frac{\gamma_d L}{U}. \tag{12b}$$

The model is closed by imposing boundary and initial conditions. These are discussed in Section 1.2.

#### 1.2 Numerical simulation of the shedding dynamics

To simulate the experimental conditions, we solve the dimensionless governing equations (Eqs. (11)) on a large, finite domain, $\hat{x} \in [0, X]$, where $X \gg 1$. Non-dimensional parameter values are discussed in Section 2 and listed in Table SM2.

##### 1.2.1 Boundary conditions

We impose no flux boundary conditions for both $\hat{C}$ and $\hat{h}$ at either side of the domain:

$$\partial_{\hat{x}} \hat{C} = 0, \quad \hat{x} \in \{0, X\}, \tag{13a}$$

$$M(\hat{h}) \frac{\partial \pi}{\partial \hat{x}} = 0, \quad \hat{x} \in \{0, X\}, \quad t > 0. \tag{13b}$$

Since the equation describing the evolution of $\hat{h}$ is fourth-order in $x$, an additional pair of boundary conditions for $\hat{h}$ is needed. We apply the natural boundary conditions:

$$\partial_{\hat{x}} \hat{h} = 0, \quad \hat{x} \in \{0, X\}, \tag{13c}$$

which correspond to vanishing contact angles at the boundaries, which are appropriate since we guarantee the droplet remains far enough from the domain boundaries.

##### 1.2.2 Initial conditions

We set the initial cell group to be a small symmetric Gaussian droplet of width $\sigma_h = \sqrt{0.1}$ centred at location $\hat{x}_0$ sufficiently far from the domain boundary (see function $\hat{h}_0$ in Figure SM3). We replicate the bacterial tracks in the 1D experiments (see Supplementary Figure 7) by considering an initially monotonically increasing bacterial profile (see Figure SM3) which saturates away from the front of the group, $\hat{B}_0(x) \approx 1$ for $\hat{x} \gg \hat{x}_0$, and decays to zero behind the group, $\hat{B}_0(\hat{x}) \approx 0$ as $\hat{x} \ll \hat{x}_0$. Without loss of generality, we equate the characteristic bacterial concentration $B_\infty$ used to scale the bacterial concentration (Eq. (10)), to the far-field bacterial concentration ahead of the cell group. We further assume that $\hat{C}_0 = \hat{B}_0$, so that it satisfies the equilibrium of the reaction term in Eq. (11b).

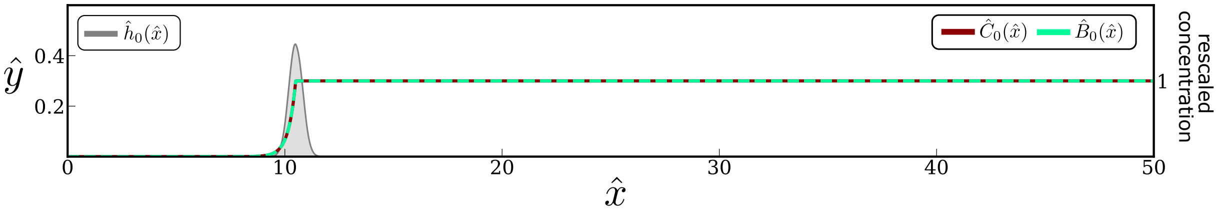


Figure SM3: Plot showing the initial conditions used in the simulations of the cell group dynamics.

##### 1.2.3 Numerical scheme

The full model requires solving three coupled non-linear partial differential equations for $\hat{h}$, $\hat{B}$ and $\hat{C}$ (see Eqs. (11)-(13)). We adopt a semi-implicit time-discretization to decouple the three equations and advance them over time (time-step size $\delta t = 0.001$). Specifically, given the approximate solution at time $t = \hat{t}_j$, $\Big(h^j(\hat{x}) = \hat{h}(\hat{t}_j, \hat{x})$, $B^j(\hat{x}) = \hat{B}(\hat{t}_j, \hat{x})$ and $C^j(\hat{x}) = \hat{C}(t_j, \hat{x})\Big)$, we proceed as follows:

1. We compute $h^{j+1}$ using Eq. (11c) and setting $\hat{C} = C^j$ and $\hat{B} = B^j$.

2. We compute $B^{j+1}$ using Eq. (11a) and setting $\hat{h} = h^{j+1}$.

3. We compute $C^{j+1}$ using Eq. (11b) and setting $\hat{B} = B^{j+1}$.

We use first-order Lagrangian finite elements to discretise the governing equations in space and divide the domain into equal intervals of size $\delta x = 0.02$. We adopt a semi-implicit time-stepping scheme to deal with the non-linar motility function $M$ in Eq. (11c), *i.e.*, we evaluate $M$ explicitly in time $M(\hat{h}) = M(h^j)$. The code is implemented in `python`, based on the `FEniCS` package for finite element methods [8]. The code is freely available at `https://github.com/giuliacelora/Dictyostelium-Group-Migration.git`.

In the numerical simulation, we adopt a smoothed version of the alignment function $S$ (see Eq. (8)):

$$S_{\alpha,\beta}(\omega) = \ln\left(2\cosh\left[\frac{\omega-\alpha}{\alpha\epsilon_-}\right]\right)\frac{\alpha\epsilon_-}{2} - \ln\left(2\cosh\left[\frac{\beta-\omega}{\beta\epsilon_+}\right]\right)\frac{\beta\epsilon_+}{2} + \frac{\alpha-\beta}{2}, \tag{14}$$

where the two hyper-parameters $0 < \epsilon_\pm \ll 1$ control the curvature of $S$ at the boundary of the linear regime, *i.e.*, $\omega = \alpha, \beta$. The results of the simulation are not sensitive to the choice of the parameters as long as they are sufficiently small. We set them to $\epsilon_+ = 0.1$ and $\epsilon_- = 0.25$. For numerical convenience, we also introduce a small level of artificial diffusion in solving the spatially-structured ODE for $\hat{B}$, with diffusion coefficient in non-dimensional units $d_b = 10^{-4}$; this is coupled to no-flux boundary conditions on $\hat{B}$ at both ends of the simulation domain.

#### 1.3 Travelling-wave model of cell group migration: Pseudo-2D active droplet

As shown in the main text, we can identify parameter regimes for which the proposed thin-film model can replicate the two-phase (travelling and shedding) cell group migration dynamics observed experimentally. The shedding of the cell group is initiated by a rapid elongation of the cell group that transitions from being a compact droplet to a multi-peaked asymmetric droplet. We are interested in understanding which mechanisms dictate this transition. We hypothesise that the elongation is driven by an imbalance between the capillary and active forces that drive the flow of cells within the droplet (away from the contact angles where the disjoining pressure becomes relevant).

To test this hypothesis, we consider a simplified version of the model for a single, self-confined droplet whose mass increases quasi-statically; this model captures all of the key features of the thin-film model presented in Section 1.1, but it is more amenable to mathematical analysis. Since we are interested in capturing solutions in which the cells are confined within a single droplet, we formulate the model into a free boundary problem, where we explicitly follow the location of the contact line, $\hat{x}_{R,F}(t)$, corresponding to the rear and front boundaries of the cell group. For simplicity, we pose the governing equations in non-dimensional form. Under the above assumptions, Eqs. (11) reduce to:

$$\frac{\partial \hat{B}}{\partial \hat{t}} = -E\hat{B}\hat{h}, \qquad \hat{x} \in \mathbb{R}, \tag{15a}$$

$$\frac{\partial \hat{C}}{\partial \hat{t}} = D_c\frac{\partial^2 \hat{C}}{\partial \hat{x}^2} - \Gamma_d\left(\hat{C} - \hat{B}\right), \qquad \hat{x} \in \mathbb{R}, \tag{15b}$$

$$\frac{\partial \hat{h}}{\partial \hat{t}} = -\frac{\partial}{\partial \hat{x}}\left(\hat{h}^2\left(\frac{\hat{h}}{3} + L_{sd}\right)\frac{\partial \pi}{\partial \hat{x}}\right) + \frac{R\hat{B}^2}{\hat{m}_B^2 + \hat{B}^2}\hat{h}, \qquad \hat{x} \in (\hat{x}_R(t), \hat{x}_F(t)), \tag{15c}$$

$$\pi = \frac{\partial^2 \hat{h}}{\partial \hat{x}^2} + Ca_\xi \hat{S}\left(\frac{\partial \hat{C}}{\partial \hat{x}}\right), \tag{15d}$$

where the cell group boundaries are implicitly defined, imposing that the droplet height first takes the value zero at these points:

$$\hat{h}(\hat{x}_R(\hat{t}), t) = \hat{h}(\hat{x}_F(\hat{t}), t) = 0, \tag{15e}$$

and the boundary conditions for the chemoattractant fields are:

$$\lim_{\hat{x}\to\infty} \hat{C}(\hat{x},\hat{t}) = 1, \quad \lim_{\hat{x}\to-\infty} \frac{\partial \hat{C}}{\partial \hat{x}}(\hat{x},\hat{t}) = 0. \tag{15f}$$

The dynamics of the floor-droplet contact points are determined by imposing conservation of mass at the moving contact lines:

$$\left.\frac{\partial \hat{h}}{\partial \hat{t}}\right|_{\hat{x}=\hat{x}_i} + \hat{x}_i'(\hat{t}) \left.\frac{\partial \hat{h}}{\partial \hat{x}}\right|_{\hat{x}=\hat{x}_i} = 0, \quad i \in \{R, F\}. \tag{15g}$$

We note that we have dropped the term associated with disjoining pressure in the definition of $\pi$ in Eq. (15d). This is because, in modelling a droplet, the contact angles are strongly imposed at the contact lines. Here, we use a dynamic contact angle model derived in [13] to capture the recession and advancement motion of the contact lines (see Figure SM4) for the moving cell group observed in the dynamic simulations. In this framework, contact angles are determined by the following boundary conditions

$$Ca_\theta \hat{x}_i'(\hat{t}) = \left.\left(\left(\frac{\partial \hat{h}}{\partial \hat{x}}\right)^2 - 1\right)\right|_{x=\hat{x}_i} n_i, \quad i \in \{R, F\}, \tag{15h}$$

where $Ca_\theta = 2U\eta_\theta/(\kappa \tan\theta_e^2)$ is the contact angle capillary number measuring the ratio between the energy dissipation at the contact lines – characterised by the friction coefficient $\eta_\theta$ – and $n_i$ indicates the outer normal at the contact lines, *i.e.*, $n_R = -1$ and $n_F = 1$. Eq. (15) is derived assuming energy is dissipated at the contact line of a moving droplet. In the case of a static droplet, the right-hand side of Eq. (15) vanishes and we recover the equilibrium contact angle condition ($\partial_{\hat{x}}\hat{h} = 1 \Rightarrow \partial_x h = \tan\theta_e$). We note that the use of Eq. (15h) is phenomenological rather than mathematically equivalent to the modulation of the contact lines in the full dynamic simulations. The model is closed by imposing appropriate initial conditions (see Section 1.2.2).

##### 1.3.1 Travelling-wave analysis.

We exploit the separation between the proliferation ($\sim$ hrs) and hydrodynamic time scales ($L/U \sim$ min) and assume that the free surface can rapidly adjust to proliferation-driven changes in the cell group volume. Hence, we adopt a quasi-steady approximation: upon small changes in the droplet mass due to proliferation, the surface $\hat{h}$, as well as the fields $\hat{B}$ and $\hat{C}$, relax to a profile that is steady in an appropriate travelling frame. The shape of the profile and the velocity of the travelling frame have to be determined as part of the solution and depends on the volume of the droplet, $\nu_{TW}$. Effectively, we therefore model the migrating cell group as a self-contained active droplet with a quasi-constant volume. We compute travelling-wave solutions by introducing the moving reference frame

$$\hat{\varphi} = \hat{x} - U_{TW}\hat{t}, \tag{16}$$

where $U_{TW}$ is the unknown velocity of the travelling-wave that we have to determine as part of our solution. We then substitute the travelling-wave ansatz

$$\hat{h}(\hat{x},\hat{t}) = h_{TW}(\hat{\varphi}), \quad \hat{B}(\hat{x},\hat{t}) = B_{TW}(\hat{\varphi}), \quad \hat{C}(\hat{x},\hat{t}) = C_{TW}(\hat{\varphi}), \tag{17}$$

into Eq. (15) to obtain:

$$U_{TW}h_{TW} = h_{TW}^2 \left[L_{sd} + \frac{h_{TW}}{3}\right] \partial_{\hat{\varphi}}\pi_{TW}, \quad \hat{\varphi} \in [0, L_{TW}], \tag{18a}$$

$$\pi_{TW} = \partial_{\hat{\varphi}\hat{\varphi}}h_{TW} + Ca_\xi S(\partial_{\hat{\varphi}}C_{TW}), \tag{18b}$$

with boundary conditions:

$$h_{TW}(0) = h_{TW}(L_{TW}) = 0, \tag{18c}$$

$$\partial_{\hat{\varphi}} h_{TW}(0) = \sqrt{1 - Ca_\theta U_{TW}}, \tag{18d}$$

$$\partial_{\hat{\varphi}} h_{TW}(L_{TW}) = -\sqrt{1 + Ca_\theta U_{TW}}. \tag{18e}$$

The length $L_{TW}$ is an unknown set by the constraining the volume of the droplet:

$$\int_0^{L_{TW}} h_{TW}(\hat{\varphi}) d\hat{\varphi} = \nu_{TW}. \tag{18f}$$

Eq. (18) appears similar to the travelling wave problem that describes the movement of passive droplets under gravity [13]. However, there are key physical and mathematical difference. Physically, active droplets move without the influence of an external field since energy is generated at the micro-scale. Mathematically, the active forcing in our model is spatially-heterogeneous and self-regulated –*i.e.*, it non-linearly depends on the travelling-wave solution via its coupling with $\partial_{\hat{\varphi}} C_{TW}$. The bacterial and chemoattract concentration profiles are determined by the following system:

$$U_{TW} \partial_{\hat{\varphi}} B_{TW} - E h_{TW} B_{TW} = 0, \tag{19a}$$

$$D_c \partial_{\hat{\varphi}\hat{\varphi}} C_{TW} + U_{TW} \partial_{\hat{\varphi}} C_{TW} + \Gamma_d (B_{TW} - C_{TW}) = 0, \tag{19b}$$

with boundary conditions:

$$\lim_{\hat{\varphi} \to \infty} B_{TW}(\hat{\varphi}) = 1, \quad \lim_{\hat{\varphi} \to -\infty} D_c \partial_{\hat{\varphi}} C_{TW} = 0, \quad \lim_{\hat{\varphi} \to \infty} C_{TW}(\hat{\varphi}) = 1. \tag{19c}$$

Eq. (19) are derived substituing Eq. (16) into Eqs. (15a)-(15b) and Eq. (15f). The far-field condition on $B_{TW}$ comes from the choice of initial conditions, which replicates the experimental 1D track experiment (see Section 1.2).

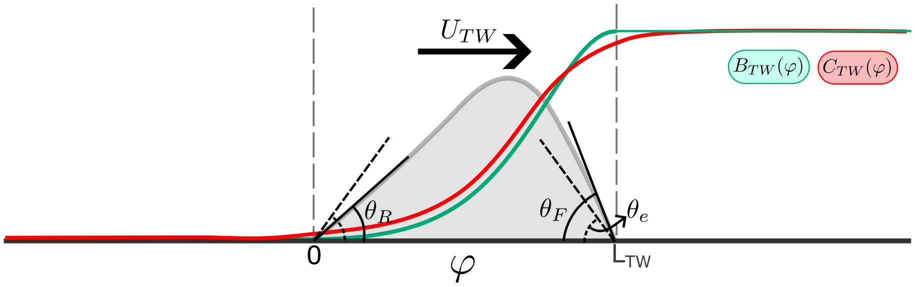


Figure SM4: Schematic illustrating how we construct the travelling wave solutions for self-contained droplet without proliferation moving at speed $U_{TW}$. Eq. (15h) is such that the front contact line will be advancing ($\theta_F > \theta_e$) while the rear contact line is receding ($\theta_F < \theta_e$).

We solve Eqs. (19) explicitly for $B_{TW}$ and $C_{TW}$ as a function of $h_{TW}$ (see Figure SM4):

$$B_{TW}(\hat{\varphi}) = \begin{cases} 1, & \hat{\varphi} > L_{TW}, \\ \exp\left[-\frac{E}{U_{TW}}\int_{\hat{\varphi}}^{L_{TW}} h_{TW}(\xi)d\xi\right], & 0 < \hat{\varphi} \le L_{TW}\ , \\ \exp\left[-\frac{E\nu_{TW}}{U_{TW}}\right], & \hat{\varphi} \le 0. \end{cases} \tag{20a}$$

$$C_{TW}(\hat{\varphi}) = \begin{cases} 1 + \left[C(L_{TW}) - 1\right] e^{\lambda_-(\hat{\varphi}-L_{TW})}, & \hat{\varphi} > L_{TW}, \\ D_c\left[\lambda_- G(\hat{\varphi}; L_{TW}) - e^{-\frac{E\nu_{TW}}{U_{TW}}}\lambda_+ G(\hat{\varphi}; 0)\right] \\ \qquad -\Gamma_d \int_0^{L_{TW}} G(\hat{\varphi}; w) B_{TW}(w) dw, & 0 \le \hat{\varphi} \le L_{TW}, \\ B_{TW}(0) + \left[C_{TW}(0) - B_{TW}(0)\right] e^{\lambda_+\hat{\varphi}}, & \hat{\varphi} < 0, \end{cases} \tag{20b}$$

where $\lambda_\pm$ are defined as:

$$\lambda_\pm = \frac{-U_{TW} \pm \sqrt{U_{TW}^2 + 4D_c\Gamma_d}}{2D_c}. \tag{20c}$$

and $G(\hat{\varphi}; \xi)$ is the Green's function associated with Eq. (19b):

$$G(\hat{\varphi}; w) = \begin{cases} \frac{1}{D_c}\frac{e^{\lambda_+(\hat{\varphi}-w)}}{\lambda_- - \lambda_+}, & 0 \le \hat{\varphi} < w, \\ \frac{1}{D_c}\frac{e^{\lambda_-(\hat{\varphi}-w)}}{\lambda_- - \lambda_+}, & w < \hat{\varphi} \le L_{TW}, \end{cases} \tag{20d}$$

Given the semi-explicit form of $B_{TW}$ and $C_{TW}$ (see Eqs. (20)), for any given value of the cell group volume $\nu_{TW}$, we solve for $h_{TW}$, $L_{TW}$ and $U_{TW}$ numerically using the Newton-Krylov method [16]. We discretize Eq. (18) using a finite volume discretisation (with a total of $N = 100$ cells). We use numerical continuation to solve the boundary value problem as a function of the cell group volume, $\nu_{TW}$. The numerical continuation is implemented in `Julia` using the `BifurcationKit.jl` for automatic bifurcation analysis [16]. The `Julia` code is freely available at `https://github.com/giuliacelora/Dictyostelium-Group-Migration.git`.

##### 1.3.2 Travelling wave results

Figure SM5 illustrates how (a) the length and (b) the velocity of travelling wave solutions depend on the cell group volume $V_{TW}$. The different curves correspond to different values of the active capillary number $Ca_\xi$. We find that for sufficiently high values of the active capillary number $Ca_\xi$, a multistability region exists whereby multiple types of travelling-wave solutions exist. The region is delimited by two fold points. This introduces the existence of a critical volume within the system (see dots in Figure SM5), $M_{cr}$, beyond which only a slow and elongated migration phenotype exists. Below the critical point (see dots in Figure SM5), capillary forces, which favour a compact cell group, dominate over active forces that favour the elongation of the cell group; hence, travelling-wave solutions are characterised by a compact profile (see Figure 3H main text). Beyond the critical point, active forces dominate over the action of surface tension; hence, travelling-wave solutions are characterised by an elongated highly asymmetric profile (see Figure 3H main text). The transition between these two regimes, which we refer to as elongation transition in the main text, occurs when active and capillary forces balance each other. When $Ca_\xi$ is small, active forces are too small to compete with the action of surface tension; hence the

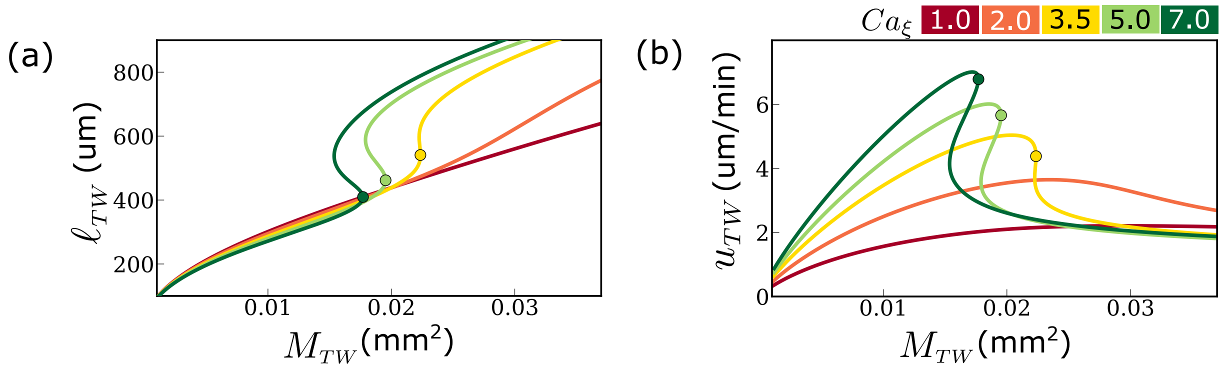


Figure SM5: The active capillary number $Ca_\xi$ controls the existence of the critical splitting mass. We plot the bifurcation diagrams of travelling-wave solutions of Eq. (18), using the cell group volume, $V_{TW} = \nu_{TW} L^2 \tan\theta_e$, as a bifurcation parameter for different values of $Ca_\xi$. We plot the (a) length and (b) velocity of the TW solution as a function of the TW volume, $V_{TW}$. The dots indicate the location of the right-most fold bifurcation when it exists. Non-dimensional parameters are set to the values in Table SM2. Note that the yellow curve corresponds to the bifurcation diagram in Figure 3G.

elongation transition is suppressed. In the dynamical simulations (see Figure 3B and Movie), as the migrating cell group crosses this critical volume it starts elongating and eventually splits. We conclude that the location and existence of the critical mass controls the shedding dynamics.

#### 1.4 Estimating flow profile for travelling wave solutions

We here discuss how we obtain the model predictions on cell velocity profiles shown in Figure 4 of the main text. Substituting the travelling-wave conditions (18a)-(18b) into Eq. (7a) and incorporating the appropriate scaling in Eq. (10)), we find that the dimensionless horizontal flow field for travelling-wave solutions is:

$$u_c^{TW}(\varphi, y) = \frac{u_{TW}}{h_{TW}(\varphi)\,(3\ell_{sd} + h_{TW}(\varphi))}\left[h_{TW}(\varphi)\,(\ell_{sd} + y) - \frac{y^2}{2}\right], \quad \varphi \in [0, \ell_{TW}], \tag{21}$$

where $u_{TW}$, $\ell_{TW} = L_{TW}$ and $\varphi = L\hat{\varphi}$ are the dimensional travelling-wave velocity, width and reference frame, respectively. Eq. (21) is used to generate the plot in Figures 4A and 4F. To generate the full flow-field in Figure 4B, we extend our lubrication analysis to approximate the leading-order expansion of the vertical component of the cell velocity $v_c^{TW}$. While this is negligible when evaluating the cell group migration, it is key to obtain the full circulatory (vortex-like) profile of cells within the cell group. In the travelling-wave analysis, proliferation is neglected and the volume of the droplet is considered to be constant. Hence the flow $\boldsymbol{u}_c^{TW}$ is incompressible, where $\boldsymbol{u}_c^{TW}(\varphi, y) = u_c^{TW}(\varphi, y)\boldsymbol{e}_x + v_c^{TW}(\varphi, y)\boldsymbol{e}_y$ (both in the Eulerian and travelling-wave reference frames). We obtain $v_c^{TW}$ by integrating the incompressibility condition and imposing a no-penetration condition at the floor ($v_c^{TW}(\varphi, 0) = 0$):

$$v_c^{TW}(\varphi, y) = \frac{3u_{TW}h_{TW}(\varphi)}{h_{TW}^2(3\ell_{sd} + h_{TW}(\varphi))^2}\left[y^3\left(\frac{\ell_{sd}}{2} + \frac{h_{TW}(\varphi)}{3}\right) - h_{TW}^2(\varphi)\left(y\,\ell_{sd} + \frac{y^2}{2}\right)\right]. \tag{22}$$

Eqs.(21)-(22) are used to generate Figure 4B, where the travelling wave solution is chosen such that $u_{TW}$ is the same as the one estimated experimentally. We note that there are two compact travelling-wave solutions (*i.e.*, both solutions belonging to the branch of the yellow curve in Figure SM5a below the critical point) that satisfy this condition due to the non-monotonic dependence of the travelling wave velocity and the volume $\nu_{TW}$. We select the travelling wave solution with the largest volume. This does not affect the qualititave nature of the velocity profile.

#### 2 Parameter values

The dimensional values of the physical parameters in the thin-film model of cell swarm migration are listed in Table SM1. We estimate most of the parameters from either the literature or experimental data (see Table SM1). The $\alpha$ and $\beta$ parameters that control the dependence of the alignment on the local chemoattractant gradients (see Eq. (14)), and the threshold concentration of bacteria for Dictyostelium cell growth $m_B$ (see Eq. (9a)) are chosen arbitrarily within the range of physically realistic values. Provided that they satisfy the physical constraints indicated in Table SM1, their precise values only have a minor influence on the numerical simulations and the inferred values of the swarm emergent material properties, *i.e.*, $\kappa/\xi$ and $\kappa/\eta$, that best capture the experimental dynamics.

The corresponding values of the non-dimensional parameters (see Eqs. (12)) adopted for the simulations of the thin-film model (Eqs. (11)) and the travelling-wave analysis (Eqs. (18) and (20)) are given in Table SM2. While the thin-film and droplet models of cell swarm migration presented in Sections 1.1 and 1.3 share most of the physical parameters, there is a difference in how the contact angle between the cell swarm and the surface is imposed. In the thin-film model, this is controlled by the disjoining pressure (Eq. (5)). In the droplet model, the contact angles are regulated by the boundary condition (15h), which depends on a single non-dimensional parameter: the contact angle capillary number $Ca_\theta$; the value of $Ca_\theta$ in Table SM2 is chosen to match the maximum value of the travelling wave velocity $U_{TW}$ (see Figure 3G main text) with the maximum value of the front velocity observed in the dynamical simulations (see Figure 3E main text).

| Parameter | Meaning | Value | Justification |
|---|---|---|---|
| $L$ | Characteristic swarm width | $100$ $[\mu m]$ | (exp) – Figure 2 main text |
| $U$ | Characteristic Dictyostelium cell speed | $12.5$ $[\mu m/min]$ | (exp) – Figure 2 main text |
| $B_\infty$ | Concentration of bacteria in the lawn | $1$ $[a.u.]$ | w.l.o.g. |
| $C_\infty$ | Concentration of chemoattractant in the lawn ($C_\infty = \gamma_p B_\infty/\gamma_d$) | $100$ $[a.u.]$ | w.l.o.g. |
| $\theta_{eq}$ | Equilibrium contact angle for shedded groups | 33 degree | (exp) – Supplementary Figure 3D |
| $\kappa/\eta$ | Ratio between emergent surface tension and emergent viscosity of the cell group | $205.4$ $[\mu m/min]$ | (exp) – Figure SM7 |
| $\kappa/\xi$ | Ratio between emergent surface tension and emergent activity of the cell group | $11.1$ $[\mu m]$ | (exp) – Figure SM7 |
| $r$ | replication rate of Dictyostelium cells growing on bacteria (doubling time 4 hr) | $0.15$ $[1/hr]$ | [5] |
| $\ell_{sd}$ | Slip length characterising friction between cell and the surface | $7.65$ $[\mu m]$ | (exp) – Figure SM6a |
| $e$ | Consumption rate of bacteria per unit swarm height | $0.0473$ $[1/(\mu m\, hr)]$ | (exp) – Figure SM6b |
| $d_c$ | Chemoattractant diffusion coefficient | $12 \times 10^3$ $[(\mu m)^2/min]$ | [15] |
| $\gamma_d$ | Decay rate of the chemoattractant | $1.2$ $[1/min]$ | decay length $L_c = \sqrt{d_c/\gamma_d}$ set to be $100\mu m$, in line with previous theoretical studies [6] |
| $h_\delta$ | Length scale at which attractive cell-floor interactions dominate | $5$ $[\mu m]$ | taken to be half the size of a cell |
| $\beta$ | Magnitude of chemoattractant gradients beyond which $S = 1$ | $2$ $[a.u./\mu m]$ | taken to be larger than the size of the chemoattractant gradients sensed by the swarm |
| $\alpha$ | Magnitude of chemoattractant gradients below which $S = 0$ | $0.02$ $[a.u./\mu m]$ | taken to be 100 times smaller than $\beta$ |
| $m_B$ | Threshold for arrest of cell proliferation | $0.05$ $[a.u.]$ | taken to be small compared to $B_\infty$ |

Table SM1: List of the dimensional physical parameters in Eqs. (1)-(9) and the values used in the simulation presented in the main text. The parameters labelled with (exp) are estimated from measured experimental data (see Section 2 for details). The highlighted parameter groupings characterise the emergent properties of the cell swarm that cannot be measured directly but are inferred to match the emergent shedding dynamics of model prediction to 1D track experiments (see Section 2.1 for more details). The remaining parameters are chosen from the literature or to satisfy realistic physical conditions (see Justification column).

We directly estimate the equilibrium contact angle $\theta_{eq}$ by measuring the contact angle formed by the shedded

| Parameter | Meaning | Value |
|---|---|---|
| $Ca_\kappa$ | Swarm capillary number | $0.2\bar{2}$ |
| $Ca_\xi$ | Swarm active capillary number | $3.5$ |
| $R$ | non-dimension proliferation rate | $0.025$ |
| $L_{sd}$ | non-dimensional slip length | $0.12$ |
| $E$ | non-dimension bacteria consumption rate | $0.51$ |
| $D_c$ | non-dimensional chemoattractant diffusion coefficient | $9.8$ |
| $\Gamma_d$ | non-dimensional chemoattractant decay rate | $9.8$ |
| $H_\delta$ | non-dimensional threshold cell-floor attraction forces | $0.0769$ |
| $\hat{\beta}$ | Magnitude of rescaled chemoattractant gradients beyond which $S = 1$ | $2$ |
| $\hat{\alpha}$ | Magnitude of rescaled chemoattractant gradients below which $S = 0$ | $0.02$ |
| $\hat{m}_B$ | Threshold for arrest cell proliferation | $0.05$ |
| $Ca_\theta$ | contact angle capillary number (chosen to match the velocity of the maximum velocity of the swarm in the TW analysis with the result from the dynamical simulations) | $0.3$ |

Table SM2: List of the non-dimensional physical parameters in the active thin-film and active droplet models.

group with the floor (see Supplementary Figure 3D). The slip length $\ell_{sd}$ is extrapolated from the flow field measured experimentally (see Figure 4C) as follows. We use Eq. (7a) to relate $\ell_{sd}$ to the ratio between the velocity at the swarm surfaces and floor, $\Delta u_c^{(i)}$, at any location, $x^{(i)}$, within the swarm; specifically

$$\ell_{sd}^{(i)} = \frac{h(x^{(i)})}{2(\Delta u_c^{(i)} - 1)}. \tag{23}$$

By sampling different locations $x^{(i)}$ within the swarm, we obtain a distribution for $\ell_{sd}^{(i)}$ (see Figure SM6a). Given the skewness of the distribution we use the median of the distribution as an estimator for the slip length (we indicate the median in Table SM1).

The rate of bacterial consumption by the swarm is estimated from the profile of bacteria quantity and swarm height in the travelling phase of the migration (Figure SM6b) using the travelling wave solution of the model (see Eq. (20a))

$$B_{exp}(x) \approx B_{exp}^0 \exp\left[-\frac{E}{V_F}\int_x^0 h_{exp}(\xi)d\xi\right] + B_{exp}^\infty. \tag{24}$$

In writing Eq. (24), we locate the origin at the position of the peak of the bacterial profile (Figure SM6b). In Eq. (24), the constant $B_{exp}^0 > 0$ accounts for the scaling of the experimental data in terms of bacteria quantity units. The constant $B_{exp}^\infty$ is introduced in the estimation of the consumption rate $E$ as a simple way to account for the non-zero value of the bacterial quantity that trails behind the Dictyostelium swarm in the experiments; we believe this is caused by complex bacterial rearrangements not accounted for in the model as they are not relevant to our main results (see Supplementary Figure 6). We fit Eq. (24) to the experimental data using `curve_fit` function in Python, which uses non-linear least square method. The estimated values of $B_{exp}^0$, $B_{exp}^\infty$ and $E$ are given in the caption of Figure SM6.

#### 2.1 Estimating the emergent material properties of the swarm

The emergent material properties of cell groups are characterised by their surface tension ($\kappa$), viscosity ($\eta$) and activity ($\xi$). We estimate the values of these parameters matching model predictions with the following metrics characterising the emergent dynamics of the swarm:

1) **time-averaged front velocity**, $V_F$ (Supplementary Figure 7B);

2) **distance between shedded groups**, $D_g$ (Supplementary Figure 7C).

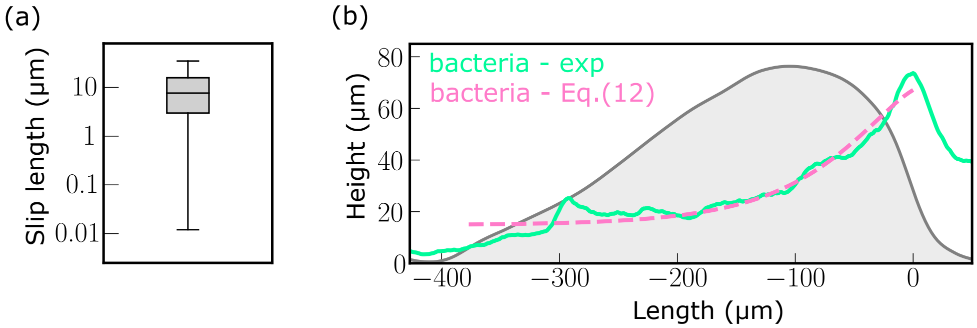


Figure SM6: (a) Estimates for the slip length ($\ell_{sd}$) applying Eq. (23) to the experimentally estimated flow field data (Figure 4 main text) – median value: $7.65\,\mu m$; interquantile range: $(2.98, 15.9)\,\mu m$. The distribution is obtained by sampling at different locations $x^{(i)}$ within the swarm. (b) Comparison of the experimental data on the bacteria quantity profile and the fitting obtained using Eq. (24) with parameters estimated via non-linear least-squares: $B^0_{exp} = 54.77\,(a.u.)$, $E = 0.51$, $B^{\infty}_{exp} = 12.4\,(a.u.)$.

Looking at Eq. (7a), the velocity field $u_c$, and therefore the emergent swarm dynamics, uniquely depend on the ratios of the emergent material properties; specifically, on $\kappa/\eta$ and $\xi/\eta$. Hence, the three parameters can not be independently identified. Therefore, we focus on inferring their relative size, which is encoded in the two capillary numbers: $Ca_\kappa$ and $Ca_\xi$ (see Eq. (12)). To estimate the values of $Ca_\kappa$ and $Ca_\xi$ from the measurements of the emergent swarm dynamics we proceed as described below.

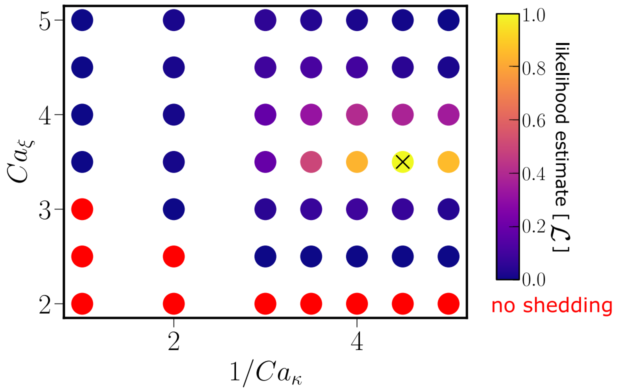


Figure SM7: Heatmap of the likelihood function $\mathcal{L}$ defined by Eq. (25); here $\mathcal{L}$ is scaled to take values in the interval $(0, 1]$. Parameters are set to the default values in Table SM2 except for the free parameters $Ca_\xi$ and $Ca_\kappa$ that we aim to estimate. We highlight in red the parameter values for which no shedding is observed for the whole duration of the simulations. We find that the likelihood is maximised for the set of capillary numbers: $Ca_\xi = 3.5$ and $Ca_\kappa^{-1} = 4.5$, for which $\chi^2 = 0.0029$, $\mathcal{L} \approx 0.997$.

Given a set of parameters ($Ca_\kappa$, $Ca_\xi$), we simulated the thin-film model (see Section 1.2) for an equivalent of 60 hours, in dimensional time units. We estimated the values of $V_F$ and $D_g$, after discarding the first 30 hours. This is to avoid the results being influenced by the choice of initial conditions. We note that the shedding of the swarm is dependent on the choice of the parameters. Parameter values that yield no shedding are discarded. If shedding is observed in the numerical simulations, we quantified the "goodness-of-fit" to the experimental data by introducing the following likelihood function:

$$\mathcal{L}(Ca_\kappa, Ca_\xi) = e^{-\chi^2(Ca_\kappa, Ca_\xi)}, \tag{25a}$$

$$\chi^2(Ca_\kappa, Ca_\xi) = \frac{1}{2}\left(\frac{V_F(Ca_\kappa, Ca_\xi) - \tilde{V}_F}{\sigma_{V_F}}\right)^2 + \frac{1}{2}\left(\frac{D_g(Ca_\kappa, Ca_\xi) - \tilde{D}_g}{\sigma_{D_g}}\right)^2. \tag{25b}$$

In Eq. (25b), the function $\chi^2$ is the standard non-linear least square functional, where $\tilde{V}_F = 4.79\,[mum/min]$ and $\sigma_F = 0.092\,[\mu m/min]$, and $\tilde{D}_g = 1.25\,[mm]$ and $\sigma_{D_g} = 0.35\,[mm]$ are the experimental mean and standard deviation of, respectively, the average front velocity and the distance between the shedded groups. The best fit is chosen as the value of $Ca_\kappa$ and $Ca_\xi$ that maximises – amongst the tested parameter values – the likelihood function $\mathcal{L}$. Results are shown in Figure SM7. We are able to identify a unique set of parameters that maximize the likelihood: $Ca_\kappa = 1/4.5$ and $Ca_\xi = 3.5$. This corresponds to the dimensional parameter groupings, $\kappa/\eta$ and $\xi/\eta$ listed in Table SM1.

We can obtain an estimate for the order of magnitude of the activity parameter $\xi$ by relating it to the strength of pulling (or traction) forces that Dictyostelium cells exert on each other, $F_{act} \approx 2 - 10 \times 10^{-8}\,[N/cell]$ [11]. Specifically, we can write $\xi = F_{act} b \rho$, where we can take $b$ to be the characteristic size of pseudopods, which we assume is comparable to the size of a cell $10\,\mu m$, and $\rho$ is directly estimated from the light sheet images (see Figure 1) $\rho = 9.4 \pm 1.1 \times 10^{-4}\,[cells/\mu m^3]$. Then we find the estimate for the activity $\xi \approx 0.17 - 1.05\,[kPa]$. This leads to the viscosities in the range $\eta \approx 0.55 - 3.4\,[kPa \cdot s]$. Our estimates for the swarm viscosity are smaller than the reported values in [2] for cell monolayers. This can be expected because of the faster dynamics of pseudopods formation/retraction compared to cell-cell adhesion. For the surface tension, we find $\kappa \approx 1.9 - 11.7\,[mN/m]$, which is within an order of magnitude of the surface tension of water ($\approx 72\,[mN/m]$).
